# Mapping the S1 and S1’ subsites of cysteine proteases with new dipeptidyl nitrile inhibitors as trypanocidal agents

**DOI:** 10.1101/760736

**Authors:** Lorenzo Cianni, Carina Lemke, Erik Gilberg, Christian Feldmann, Fabiana Rosini, Fernanda dos Reis Rocho, Jean F.R. Ribeiro, Daiane Y. Tezuka, Carla D. Lopes, Sérgio de Albuquerque, Jürgen Bajorath, Stefan Laufer, Andrei Leitão, Michael Gütschow, Carlos A. Montanari

## Abstract

The cysteine protease cruzipain is considered to be a validated target for therapeutic intervention in the treatment of Chagas disease. A series of 26 new compounds waswere designed, synthesized, and tested against the recombinant cruzain (Cz) to map its S1/S1′ subsites. The same series was evaluated on a panel of four human cysteine proteases (CatB, CatK, CatL, CatS) and *Leishmania mexicana* CPB, which is a potential target for the treatment of cutaneous leishmaniasis. The synthesized compounds are dipeptidyl nitriles designed based on the most promising combinations of different moieties in P1 (ten), P2 (six), and P3 (four different building blocks). Eight compounds exhibited a *K*_i_ smaller than 20.0 nM for Cz, whereas three compounds met these criteria for LmCPB. The three inhibitors had an EC_50_ value of ca. 4 μM, thus being equipotent to benznidazole according to the anti-trypanosomal effects. Our mapping approach and the respective structure-activity relationships provide insights into the specific ligand-target interactions for therapeutically relevant cysteine proteases.

**Author Summary:** Despite many achievements in identifying novel agents for the treatment of tropical and neglected diseases, further research continues to be of fundamental importance. Our research groups have been using the cruzipain cysteine protease in its recombinant form, cruzain (Cz), to identify new trypanocidal agents. Considering the possible interchangeability with other cysteine proteases, the same series of dipeptidyl nitriles was tested in *Leishmania mexicana* LmCPB. Other potential targets for such inhibitors are human cysteine cathepsins, which are involved in different disease states. Thus, the inhibitors were also tested against cathepsins B, L, K, and S. Our results demonstrate that inhibition of these cysteine proteases can be achieved by appropriate structural modifications of dipeptidyl nitriles. It was also possible to identify trypanocidal agents, equipotent to benznidazole, the current drug of choice used for the treatment of Chagas disease.

## Introduction

Chagas disease, aka American trypanosomiasis, is a serious health and social problem in Latin America and new non-endemic areas such as Japan, East Europe, and the USA. Chagas disease has an annual incidence of 30,000 new cases and 14,000 deaths per year. In addition, more than 70,000 million people living in areas where they are at risk of contracting the disease [1].

The etiological agent, the protozoa parasite *Trypanosoma cruzi* (*T. cruzi*), is transmitted by blood-sucking reduviid bugs of the subfamily Triatominae [2]. The only two existing drugs in the market, benznidazole and nifurtimox, show strong side effects and inefficiency in the chronic stage of the disease [3, 4]. New safe and efficacious drugs are therefore required to address with these still unmet medical needs. Initiatives such as the one launched by the Drugs for Neglected Diseases (DNDi) have led to worldwide collaborative efforts to discover new therapeutic targets [5]. Cruzain (Cz), a recombinant form of the enzyme cruzipain (EC 3.4.22.51) [6], is the most abundant cysteine protease (CP) present in the parasite and essential for its development and survival inside and outside the host cell in all forms of its life cycle. This makes Cz a druggable target for the development of new chemotherapeutic agents against Chagas disease [7, 8].

Cz represents a target for irreversible (or suicide) and reversible inhibitors. K777 was at the forefront of the first generation of irreversible Cz inhibitors and initially characterized by the Sandler Center for Research in Tropical Parasitic Disease (University of California, San Francisco) [9]. Despite its ability to rescue mice of a lethal experimental *T*. c*ruzi* infection and reduce parasite growth in dogs, preclinical safety and toxicology studies revealed substantial side effects of K777 in primates and dogs, even when administered in low doses [10, 11]. Current research is being focused on reversible Cz inhibitors as these are assumed to overcome possible off-target effects [12].

The structure of Cz is closely related to those of mammalian CPs (CatL, CatK and CatS). Three-dimensional (3D) structures of Cz a variety of ligands have already been resolved [13], enabling the applicability of target-based molecular design to find Cz-inhibiting P1, P2, and P3 positions of dipeptidyl nitrile ligands with the respective subsites of the enzyme (S1, S2, S3) and the trypanocidal activities of such inhibitors [14]. In this study, we designed a new, structurally expanded series of 26 Cz-inhibiting dipeptidyl nitriles, in particular by leveraging the P1-S1/S1′ interactions. We explored the structure-activity relationships (SARs), mapped the active site of the target enzyme and evaluated the antichagasic properties of the compounds. Besides that, we have tested them against four human cysteine cathepsins (CatB, CatL, CatK, CatS) all of which constituting important targets for human diseases [15], and against the cysteine protease LmCPB, a novel macromolecular target to fight *Leishmania mexicana.* As a result of this study, several new low-nanomolar inhibitors of different CPs were discovered and the action of three representatives on *T. cruzi*, being equipotent to benznidazole, was characterized.

## Methods

### Modelling

Putative binding modes of novel dipeptidyl nitrile inhibitors compounds were derived from the crystal structure of *N*-(2-aminoethyl)-alpha-benzoyl-l-phenylalaninamide (33L) bound to Cz (PDB ID: 4QH6). This ligand-target-complex served as a template for knowledge-based modelling and was preprocessed using the “Structure Preparation” and “Protonate3D”-tools of the modeling software “Molecular Operating Environment” (MOE) [16], version 2018.0101, with default settings. By modification of moieties, the cocrystallized ligand was structurally transformed to the compound of interest. Obtained conformations were optimized using the force field “Amber10:EHT”.

### Synthetic chemistry

#### General considerations

Synthesis was performed as summarized in Fig 1 and Fig 2. Melting points were determined on a Büchi 510 oil bath apparatus and are uncorrected. Infrared spectra were obtained from FT-IR Thermo Scientific Nicolet 380. Reagents, starting materials and solvents were of commercial quality and were used without further purification unless otherwise stated. All syntheses were started with enantiopure amino acids. TLC analysis was carried out on Merck 60 F_254_ silica gel plates and visualized under UV light at 254 nm and 365 nm or by using ninhydrin staining solution. Preparative column chromatography was carried out on Grace Davison Davisil LC60A 20-45 micron or Merck Geduran Si60 63-200 micron silica using the Interchim PuriFlash 430 automated flash chromatography system. The purity of all tested compounds was determined with one of the three protocols (A-C) noted below.

Purity was determined via RP-HPLC on a Hewlett Packard 1090 Series II LC with a Phenomenex Luna C18 column (150 × 4.6 mm, 5 μm) and detection was performed by a UV DAD (200 – 440 nm). Elution was carried out with the following gradient: 0.01 M KH_2_PO_4_, pH 2.30 (solvent A), MeOH (solvent B), 40% B to 85% B in 8 min, 85% B for 5 min, 85% to 40 % B in 1 min, 40% B for 2 min, stop time 16 min, flow 1.5 mL/min.
Purity was determined using an LC-MS instrument (ABSCIEX API 2000 LC-MS/MS, HPLC Agilent 1100) with a Phenomenex Luna C18 HPLC column (50 × 2.00 mm, 3 µm) and detection was performed by a UV DAD (200 – 440 nm). Elution was carried out with the following gradient: 0.02 M NH_4_CH_3_CO_2_, pH 7.0 (solvent A), MeOH (solvent B) start with 100%, 10% B in 20 min to 100% B, 10 min 100% B, stop time 20 min, flow 0.25 mL/min.
Purity was determined with an LC-MS instrument (AmaZon SL ESI-MS, Shimadzu LC) with a cellulose-2 Phenomenex column (250 × 4.6 mm, 5 μm) or a Diacel column (IC-chiralpak, 250 × 4.6 mm, 5 μm). An isocratic elution with MeCN and water was applied as specified, stop time 60 min, flow 0.5 mL/min.

**Fig 1.**
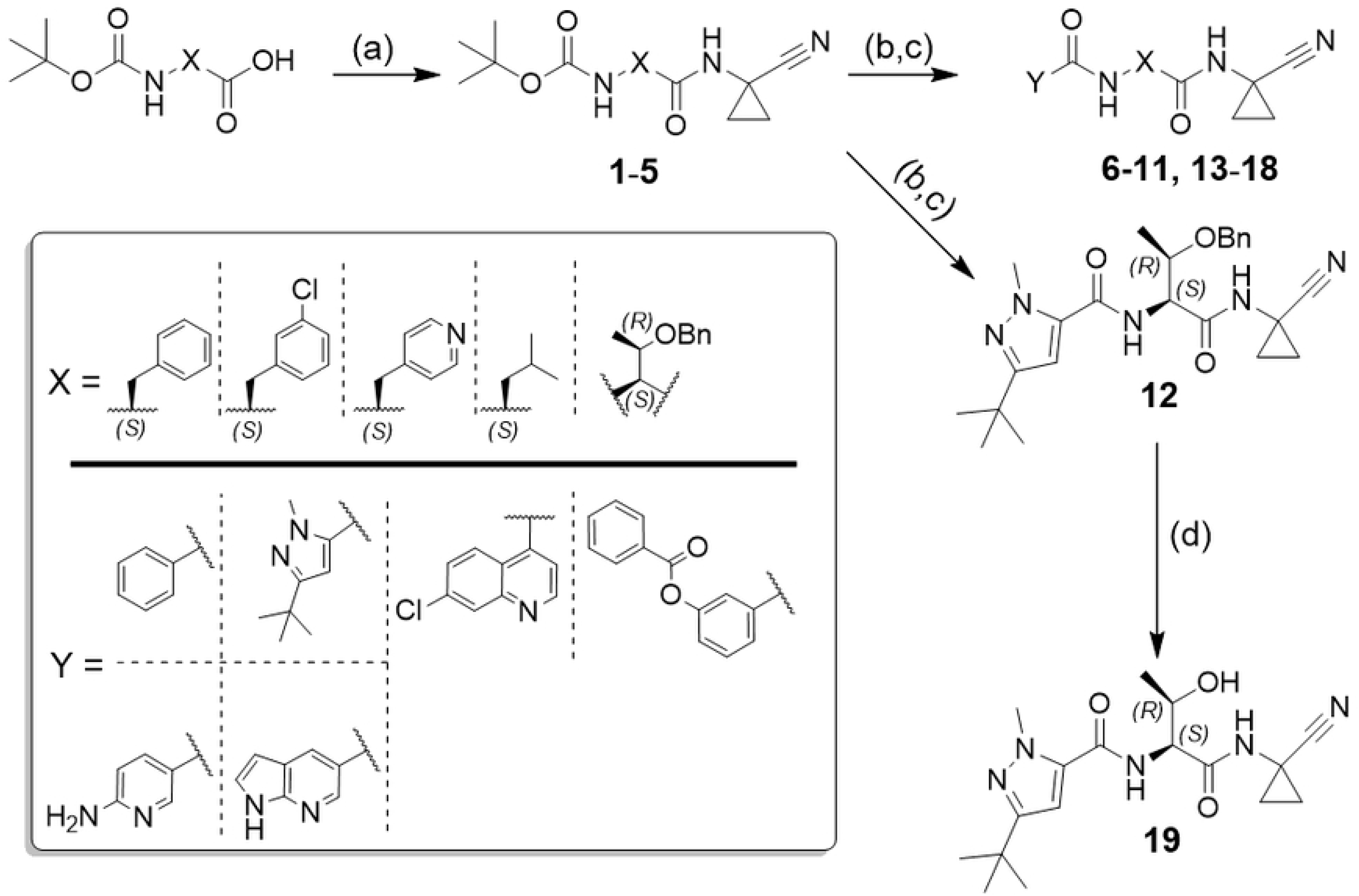
General synthesis of compounds 6-19. Reagents and conditions: a) HATU, DIPEA, 1-amino-1-cyclopropanecarbonitrile, DMF, rt, 18 h; b) formic acid, rt, 18 h; c) HATU or TBTU, DIPEA, carboxylic acid, DMF, rt, 18 h; d) DDQ, DCM, rt, 18 h.

**Fig 2.**
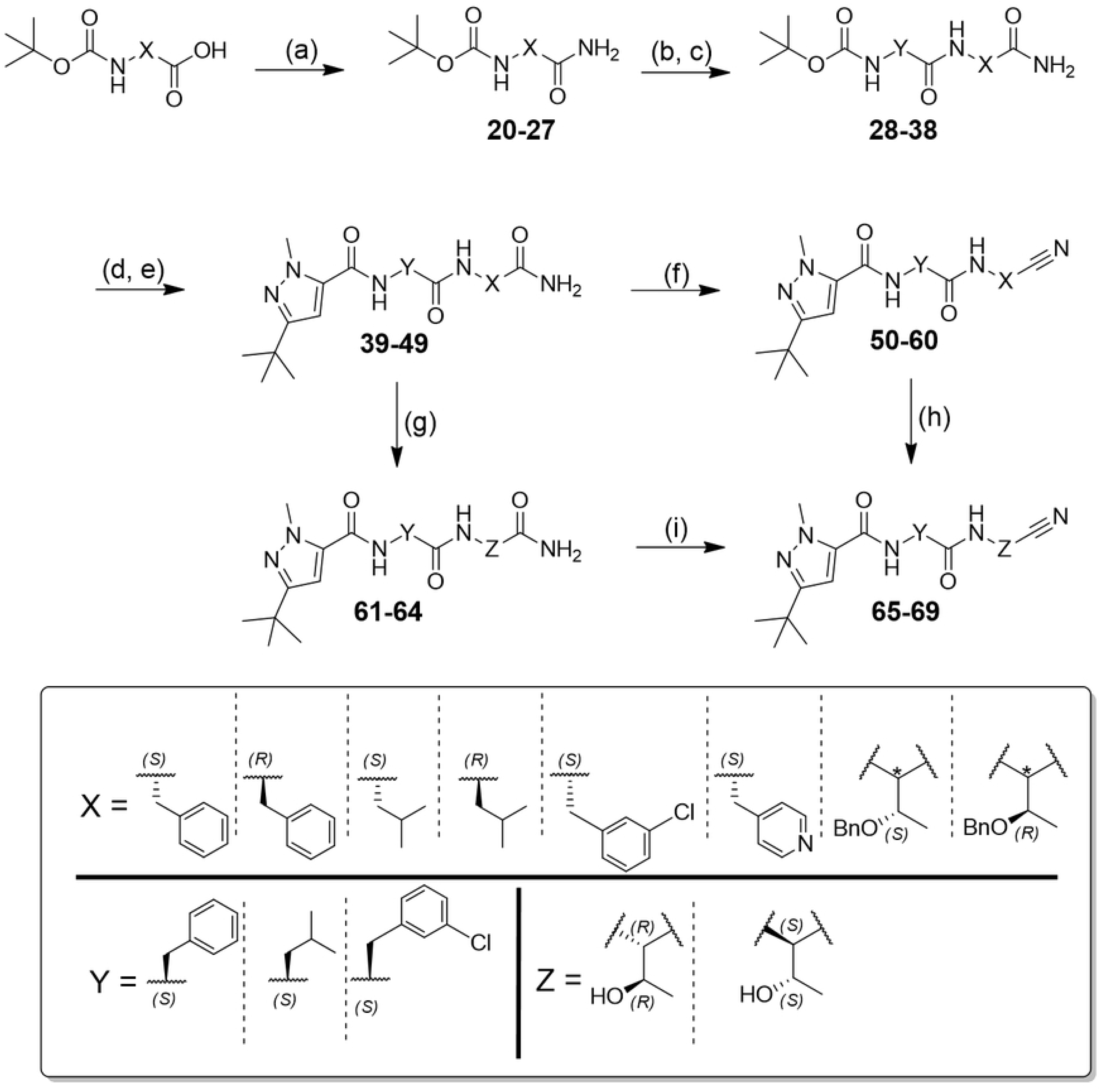
General synthesis of compounds 50-60, 65-69. Reagents and conditions: a) Isobutyl chloroformate, NH_4_Cl 2 M, DIPEA, DMF, 0 °C to rt, 20 h; b) TFA, CH_2_Cl_2_, 0 °C to rt, 2 h; c) HATU, DIPEA, Boc-AA-OH, DMF, rt, 18 h; d) TFA, CH_2_Cl_2_, 0 °C to rt, 2 h; e)TBTU, DIPEA, 3-(*tert*-butyl)-1-methyl-1*H*-pyrazole-5-carboxylic acid, DMF/CH_2_Cl_2_, rt, 18 h; f) Cyanuric chloride, DMF, 0 °C to rt, 0.5 h; g) H_2_ (1 atm), Pd/C, rt, 18 h; h) *p*-TsCl, Py, rt, 3-5 days; i) TFAA, DIPEA, THF, 0 °C to rt, 2 h.

NMR spectra were recorded on Bruker Avance 200 MHz, Bruker Avance 400 MHz, and Bruker Avance DRX 500 MHz NMR spectrometers. Chemical shifts are reported in ppm relative to TMS or the residual proton peak of the re-protonated deuterated solvent, and the spectra were calibrated against the residual proton peak of the used deuterated solvent. The following symbols indicate spin multiplicities: s (singlet), s br (broad singlet), d (doublet), dd (doublet of doublet), t (triplet), tt (triplet of triplet), q (quartet), sept (septet), and m (multiplet). Standard mass spectra were obtained either as ESI-MS (pos. and/or neg. mode) from a Advion DCMS interface, (settings as follows: ESI voltage 3,50 kV, capillary voltage 187 V, source voltage 44 V, capillary temperature 250 °C, desolvation gas temperature 250 °C, gas flow 5 L/min) or by an API 2000 mass spectrometer (electron spray ion source, ABSCIEX, Darmstadt, Germany) coupled to an Agilent 1100 HPLC system.

HRMS spectra were recorded on a Bruker micrOTOF-Q mass spectrometer connected to a Thermo Scientific Dionex UltiMate 3000 LC via an ESI interface using a Nucleodur C18 Gravity column (50 × 2.0 mm, 3 μm) or were recorded on Thermo Scientific LTQ Velos Orbitrap, in electrospray ionization (ESI) mode by direct injection.

The synthetic route was developed to optimize the set of substituents to be placed in P1, P2, and P3 that have been defined after the planning and design studies. Due to the diversity of building blocks, it was necessary to evaluate different coupling and dehydrating reagents, aiming at the best yield and preventing racemization.**Error! Bookmark not defined.**

#### General procedure for amide synthesis

**Method A:** Isobutyl chloroformate (790 mg, 0.75 mL, 5.5 mmol, 1.1 equiv) was added dropwise to a solution of Boc-(*R* or *S*) amino acid (5.0 mmol, 1.0 equiv.), DIPEA (1.6 g, 2.28 mL, 13.0 mmol, 2.6 equiv.) in dry DMF (20 mL), under argon atmosphere, at −30 °C and it was stirred for 0.5 h. Then, an aqueous 2 M NH_4_Cl solution (294 mg, 2.75 mL, 5.50 mmol, 1.1 equiv.) was added. The resulting solution was stirred at room temperature for 20 h. The reaction mixture was dried under reduced pressure. Ethyl acetate (100 mL) was added, and it was washed with a saturated NaHCO_3_ solution (3 × 50 mL) and brine (1 × 50 mL). The organic phase was dried over Na_2_SO_4_ and evaporated under reduced pressure to give a crude residue that was purified by flash column chromatography.

**Method B:** The free primary amine (1.0 mmol, 1.0 equiv.) was added to a solution of the carboxylic acid (1.3 mmol, 1.3 equiv.), HATU (490 mg, 1.3 mmol, 1.3 equiv.) and DIPEA (364 mg, 0.45 mL, 2.60 mmol, 2.6 equiv.) in dry DMF (5 mL) under argon atmosphere. The resulting solution was stirred at room temperature for 20 h. The reaction mixture was diluted with ethyl acetate (10 mL) and washed with a saturated NaHCO_3_ solution (3 × 20 mL) and brine (3 × 20 mL). The organic phase was dried over Na_2_SO_4_ and evaporated to give a crude residue that was purified by flash column chromatography.

**Method C:** The free primary amine (1.0 mmol, 1.0 equiv.) was added to a solution of the carboxylic acid (1.3 mmol, 1.3 equiv.), TBTU (410 mg, 1.30 mmol, 1.3 equiv.) and DIPEA (364 mg, 0.45 mL, 2.60 mmol) in dry DMF/CH_2_Cl_2_ (1:1, 10 mL) under argon atmosphere. The resulting solution was stirred at room temperature for 20 h. The reaction mixture was diluted with ethyl acetate (10 mL) and washed with a saturated NaHCO_3_ solution (3 × 20 mL) and brine (3 × 20 mL). The organic phase was dried over Na_2_SO_4_ and evaporated to give a crude residue that was purified by flash column chromatography.

#### General procedure for removal of the Boc protecting group

**Method A:** The Boc-protected amino compound (0.25 mmol, 1.0 equiv.) was treated with formic acid (2.44 g, 2.0 mL, 47.9 mmol, 47.9 equiv.) at room temperature. The resulting solution was stirred for 18 h. The reaction mixture was evaporated under reduced pressure to get a yellowish oil. It was treated with an aqueous solution of 1.0 M NaOH until pH 9 was reached. The product was extracted with ethyl acetate (4 × 20 mL) and then washed with brine (1 × 20 mL). The organic phase was evaporated to obtain a colorless oil. The formation of the product was confirmed by TLC (ethyl acetate). The product was used for the next step without further purification.

**Method B:** To a solution of Boc-protected amino compound (1.0 mmol, 1.0 equiv.) in dry CH_2_Cl_2_ (3 mL) was added TFA (912 mg, 0.91 mL, 8.00 mmol, 8.0 equiv.) at 0 °C. The mixture was stirred and allowed to reach room temperature within 2 h. The progress of the reaction was monitored by TLC (ethyl acetate). The reaction mixture was evaporated under reduced pressure to eliminate the excess of TFA to get a yellowish solid. The product was used for the next step without further purification.

#### General procedure for dehydration of primary amides to nitriles

**Method A:** The primary amide (1.0 mmol, 1.0 equiv.) was dissolved in dry DMF (5 mL) at 0 °C. Then, cyanuric chloride (73 mg, 0.4 mmol, 1.1 equiv.) was slowly added to the solution under argon atmosphere. The resulted solution was stirred for 0.5 h. Saturated NaHCO_3_ solution (30 mL) was added and it was stirred at room temperature for 2 h. The product was extracted with ethyl acetate (2 × 50 mL), and then the reunited organic phases were washed with an aqueous solution of 1.0 M KHSO_4_ (3 × 20 mL), brine (4 × 30 mL) and dried over Na_2_SO_4_. The solvent was removed, and the crude product was purified by flash silica gel chromatography.

**Method B:** The primary amide (1.0 mmol, 1.0 equiv.) was dissolved in dry THF (5 mL) and DIPEA (364 mg, 0.45 mL, 2.6 mmol, 2.6 equiv.) was added. Trifluoroacetic anhydride (273 mg, 0.18 mL, 1.30 mmol, 1.3 equiv.), was added over 5 min, at 0 °C. The mixture was stirred and allowed to reach room temperature within 2 h. Then the reaction was quenched with H_2_O (20 mL), THF removed in vacuo, and the product was extracted into ethyl acetate (2 × 50 mL). The organic phase was washed with a solution 1.0 M of KHSO_4_ (3 × 20 mL) and with a saturated NaHCO_3_ solution (3 × 20 mL) and brine (3 × 20 mL) and dried over Na_2_SO_4_. The solvent was removed, and the crude product was purified by flash silica gel chromatography.

**Method C:** The primary amide (1.0 mmol, 1.0 equiv.) was dissolved in dry pyridine (5 mL) at room temperature. Then, *p*-toluenesulfonyl chloride (572 mg, 3.0 mmol, 3.0 equiv.) was added to the solution under argon atmosphere. The resulting solution was stirred for 3 days. Upon addition of saturated NaHCO_3_ solution (30 mL), the reaction mixture was stirred at room temperature for 2 h. The solution was dried under reduced pressure. The product was extracted with ethyl acetate (2 × 50 mL), and then the reunited organic phases were washed with a 1.0 M solution of KHSO_4_ (2 × 20 mL), brine (4 × 30 mL) and dried over Na_2_SO_4_. The solvent was removed, and the crude product was purified by flash silica gel chromatography.

#### General procedure for removal of the benzyl protecting group

**Method A:** The corresponding protected threonine (1.0 mmol, 1.0 equiv.) was dissolved in ethanol absolute (20 mL) in an argon atmosphere. Upon addition of 10% Pd/C, H_2_ was bubbled in the solution for 0.5 h. The resulting solution was stirred under H_2_ atmosphere for 12 h. The progress of the reaction was monitored by TLC (ethyl acetate). The solution was filtered on celite two times and dried under reduced pressure to afford the desired product as a colorless wax. The product was used for the next step without further purification.

**Method B:** The corresponding protected threonine (1.0 mmol, 1.0 equiv.) was dissolved in dry CH_2_Cl_2_ (20 mL) under argon atmosphere. Then, DDQ (908 mg, 4.0 mmol, 4.0 equiv.) was added, and the resulting solution was stirred for 4 days at room temperature. The progress of the reaction was monitored by TLC (ethyl acetate). The reaction was quenched with an aqueous 1.0 M solution of NaHSO_3_ (20 mL). Then, CH_2_Cl_2_ was removed under reduced pressure. The product was extracted with ethyl acetate (2 × 50 mL), and the reunited organic phases were washed with an aqueous solution of 1.0 M KHSO_4_ (2 × 20 mL), brine (4 × 30 mL) and dried over Na_2_SO_4_. The solvent was removed, and the crude product was purified by flash silica gel chromatography.

#### Synthesis and characterization of compound 1-5

Compounds **1**-**5** have been synthesized from the corresponding amino acid and 1-amino-1-cyclopropanecarbonitrile following the general procedure for amide synthesis (method B) [14].

*(S)-tert-Butyl (1-((1-cyanocyclopropyl)amino)-1-oxo-3-phenylpropan-2-yl)carbamate (**1**)*

Yield 92%. White solid. *R_f_* = 0.9 (ethyl acetate: *n*-hexane; 7:3). Mp. 146–147 °C. ^1^H NMR (500 MHz, CDCl_3_) δ 7.27 – 7.04 (m, 5H), 4.58 (m, 1H), 3.19 (dd, *J* = 13.8, 5.0 Hz, 1H), 2.88 (dd, *J* = 9.5, 5.0 Hz, 1H), 1.31 (s, 9H), 1.25 (m, 2H), 1.04 (m, 2H). ^13^C NMR. (125 MHz, CDCl_3_) δ 173.06, 155.30, 137.73, 129.35, 128.19, 126.45, 120.80, 78.30, 55.54, 37.36, 28.26, 19.77, 15.80, 15.75. ESI-MS (+) Calc. for [C_18_H_23_N_3_O_3_] 329.39, found: 352.3 [M+Na]^+^.

*(S)-tert-Butyl (3-(3-chlorophenyl)-1-((1-cyanocyclopropyl)amino)-1-oxopropan-2-yl)carbamate (**2**)*

Yield 83%. White solid. *R_f_* = 0.7 (ethyl acetate: *n*-hexane; 6:4). Mp. 146–147 °C. ^1^H NMR (500 MHz, CDCl_3_) δ 7.26 – 7.25 (m, 1H), 7.23 – 7.21 (m, 1H), 7.08 – 7.03 (m, 2H), 4.27 – 4.24 (m, 1H), 3.10 (dd, *J* = 13.8, 5.0 Hz, 1H), 2.83 (dd, *J* = 9.5, 5.0 Hz, 1H), 1.52 – 1.44 (m, 2H), 1.41 (s, 9H), 1.13 – 1.05 (m, 2H). ^13^C NMR (125 MHz, CDCl_3_) 172.02, 155.61, 138.23, 134.32, 129.29, 127.49, 127.25, 119.48, 119.48, 80.55, 55.28, 37.87, 28.18, 20.14, 16.68, 16.56. ESI-MS (+) Calc. for [C_18_H_22_ClN_3_O_3_] 363.83, found: 364.3 [M+H]^+^.

*(S)-tert-Butyl (1-((1-cyanocyclopropyl)amino)-1-oxo-3-(pyridin-4-yl)propan-2-yl)carbamate (**3**)*

Yield 75%. White solid. *R_f_* = 0.5 (ethyl acetate: *n*-hexane; 4:6). Mp. 134–135 °C. ^1^H NMR (200 MHz, CD_3_OD) δ 8.45 (d, *J* = 4.9 Hz, 2H), 7.34 (d, *J* = 5.8 Hz, 2H), 4.36 – 4.23 (m, 1H), 2.96 (dd, *J* = 14.0, 5.0 Hz, 2H), 1.53 – 1.46 (m, 1H), 1.38 (s, 9H), 1.20 – 1.14 (m, 2H). ^13^C NMR (50 MHz, CDCl_3_) δ 174.74, 149.95, 149.15, 126.52, 121.09, 80.79, 56.03, 38.88, 38.31, 28.55, 21.21, 17.05. ESI-MS (+) Calc. for [C_17_H_22_N_4_O_3_] 330.38, found: 331.2 [M+H]^+^.

*(S)-tert-Butyl (1-((1-cyanocyclopropyl)amino)-4-methyl-1-oxopentan-2-yl)carbamate (**4**)*

Yield 61%. White solid. *R_f_* = 0.7 (ethyl acetate: *n*-hexane; 4:6). Mp. 162–164 °C. ^1^H NMR (500 MHz, DMSO-*d_6_*) δ 8.76 (s, 1H), 6.86 (d, *J* = 7.9 Hz, 1H), 3.86 (dt, *J* = 8.7, 5.5 Hz, 1H), 1.55-1.54 (m, *J* = 6.6 Hz, 1H), 1.44 (dd, *J* = 7.9, 5.5 Hz, 2H), 1.37 – 1.42 (m, 2H), 1.36 (s, 9H), 1.07 (dd, *J* = 7.7, 5.3 Hz, 2H), 0.84 (2d, *J* = 6.6 Hz, 6H). ^13^C NMR (125 MHz, DMSO-*d_6_*) δ 174.17, 155.50, 120.92, 78.22, 52.58, 40.49, 28.31, 24.38, 23.01, 21.66, 19.91, 15.87, 15.75. ESI-MS (+) Calc. for [C_15_H_25_N_3_O_3_] 295.37, found: 318.3 [M+Na]^+^.

*tert-Butyl ((2S,3R)-3-(benzyloxy)-1-((1-cyanocyclopropyl)amino)-1-oxobutan-2-yl)carbamate (**5**)*

Yield 89%. White solid. *R_f_* = 0.7 (ethyl acetate: *n*-hexane; 4:6). Mp. 88–90 °C. ^1^H NMR (200 MHz, CDCl_3_) δ 7.36 – 7.32 (m, 5H), 4.78 – 4.60 (m, 3H), 4.22 – 4.12 (m, 1H), 1.57 – 1.44 (m, 2H), 1.30 – 1.27 (m, 9H), 1.17 – 1.13 (m, 5H). ESI-MS (+) Calc. for [C_20_H_27_N_3_O_4_] 373.44, found: 396.4 [M+Na]^+^.

#### Synthesis of compounds 6-18

Compounds **6**-**18** have been synthesized in two steps from compounds **1**-**5**. First, the Boc group was removed (procedure A), and then the free amine was coupled to the carboxylic acid following the general procedure for amide synthesis (method B or method C, as indicated).

Synthesis and characterization of compounds **6**, **9** and **11** have been already published elsewhere [13].

*(S)-N-(3-(3-Chlorophenyl)-1-((1-cyanocyclopropyl)amino)-1-oxopropan-2-yl)benzamide (**7**)*

Method B. Yield 86%. White solid. *R_f_* = 0.7 (ethyl acetate: *n*-hexane; 6:4). Mp. 213 – 215 °C. ^1^H NMR (500 MHz, DMSO-*d_6_*) δ 9.04 (s, 1H), 8.67 (d, *J* = 8.1 Hz, 1H), 7.84 – 7.83 (m, 2H), 7.56 – 7.54 (m, 1H), 7.53 (t, *J* = 7.7 Hz, 2H), 7.41 (s, 1H), 7.32 – 7.25 (m, 3H), 4.65 – 4.60 (m, 1H), 3.09 (dd, *J* = 13.6, 5.0 Hz, 1H), 3.02 (dd, *J* = 15.2, 5.0 Hz, 1H), 1.51 – 1.49 (m, 2H), 1.12 – 1.06 (m, 2H). ^13^C NMR (125 MHz, DMSO-*d_6_*) δ 172.57, 166.48, 140.62, 132.82, 131.52, 130.04, 129.18, 128.31, 128.05, 127.58, 126.50, 120.81, 54.47, 36.60, 19.90, 15.82. HRMS (+) Calc. for [C_20_H_19_ClN_3_O_2_]^+^ 368.11658, found: 368.11615 [M+H]^+^. HPLC (protocol B): t_R_ (min) = 10.29. Purity: 99.6%.

*(S)-3-(tert-Butyl)-N-(3-(3-chlorophenyl)-1-((1-cyanocyclopropyl)amino)-1-oxopropan-2-yl)-1-methyl-1H-pyrazole-5-carboxamide (**8**)*

Method C. Method B. Yield 72%. Yellowish solid. *R_f_* = 0.7 (ethyl acetate: *n*-hexane; 5:5). Mp. 152 – 154 °C. ^1^H NMR (400 MHz, DMSO-*d_6_*) δ 9.03 (s, 1H), 8.59 (d, *J* = 8.4 Hz, 1H), 7.39 (s, 1H), 7.31 – 7.24 (m, 3H), 6.79 (s, 1H), 4.57 – 4.53 (m, 1H), 3.88 (s, 3H), 3.08 (dd, *J* = 13.6, 5.1 Hz, 1H), 2.94 (dd, *J* = 13.6, 10.2 Hz, 1H), 1.50 – 1.47 (m, 2H), 1.28 (s, 9H), 1.10 – 1.05 (m, 2H). ^13^C NMR (100 MHz, DMSO-*d_6_*) δ 172.10, 159.46, 158.70, 140.26, 134.95, 132.64, 129.94, 129.15, 127.85, 126.39, 120.63, 103.82, 55.21, 36.38, 31.58, 30.36, 19.75, 15.67, 15.61. HRMS (+) Calc. for [C_22_H_27_ClN_5_O_2_]^+^ 428.18533, found: 428.1864 [M+H]^+^. HPLC (protocol B): t_R_ (min)= 11.17. Purity: 98.3%.

*(S)-3-(tert-Butyl)-N-(1-((1-cyanocyclopropyl)amino)-1-oxo-3-(pyridin-4-yl)propan-2-yl)-1-methyl-1H-pyrazole-5-carboxamide (**10**)*

Method C. Yield 56%. Yellowish oil. *R_f_* = 0.4 (ethyl acetate: *n*-hexane; 5:5). ^1^H NMR (200 MHz, CD_3_OD) δ 8.48 – 8.46 (m, 2H), 7.43 (d, *J* = 4.9 Hz, 2H), 6.69 (s, 1H), 4.86 – 4.78 (m, 1H), 3.95 (s, 3H), 3.04 – 3.02 (m, 1H), 2.89 – 2.84 (m, 1H), 1.56 – 1.50 (m, 2H), 1.31 (d, *J* = 4.3 Hz, 9H), 1.22 – 1.18 (m, 2H). ^13^C NMR (50 MHz, CD_3_OD) δ 172.80, 160.35, 160.05, 148.62, 147.59, 134.80, 125.00, 119.66, 103.60, 53.18, 37.37, 36.45, 31.42, 29.44, 19.92, 15.60, 15.28. FT-IR (KBr, cm^−1^) 3297.16, 2966.17, 2242.31, 1670.63, 1601.36, 1531.90, 1425.65, 1352.10, 1278.55, 1241.77, 1049.72, 988.42, 808.63, 755.51, 722.82, 511.19, 506.24, 489.90. HRMS (+) Calc. for [C_21_H_27_N_6_O_2_]^+^ 395.21955, found: 395.21973 [M+H]^+^. HPLC (protocol B): t_R_ (min) = 3.68. Purity: 94.7%.

*N-((2S,3R)-3-(Benzyloxy)-1-((1-cyanocyclopropyl)amino)-1-oxobutan-2-yl)-3-(tert-butyl)-1-methyl-1H-pyrazole-5-carboxamide (**12**)*

Method C. Yield 53%. Yellowish solid. *R_f_* = 0.4 (ethyl acetate: *n*-hexane; 6:4). Mp. 100 – 101 °C. ^1^H NMR (200 MHz, CDCl_3_) δ 7.36 – 7.32 (m, 5H), 6.45 (s, 1H), 4.76 – 4.62 (m, 3H), 4.22 – 4.12 (m, 1H), 4.09 (s, 3H), 1.57 – 1.44 (m, 2H), 1.30 – 1.27 (m, 9H), 1.20 – 1.13 (m, 5H). ^13^C NMR (50 MHz, CDCl_3_) δ 170.36, 160.18, 159.74, 137.15, 134.06, 128.48, 127.98, 127.68, 119.25, 103.18, 74.03, 71.48, 54.81, 38.38, 31.66, 30.16, 28.68, 20.24, 16.91, 15.85. FT-IR (KBr, cm^−1^) 3288.99, 2953.92, 2917.14, 2214.31, 1646.31, 1589.10, 1495.12, 1049.31, 1294.89, 1200.91, 1033.37, 922.51, 755.51, 681.95. HRMS (+) Calc. for [C_24_H_32_N_5_O_3_]^+^ 437.25051, found: 438.25102 [M+H]^+^. HPLC (protocol B): t_R_ (min) = 8.77. Purity: 97.0%.

*(S)-7-Chloro-N-(1-((1-cyanocyclopropyl)amino)-1-oxo-3-phenylpropan-2-yl)quinoline-4-carboxamide (**13**)*

Method B. Yield 74%. White solid. *R_f_* = 0.7 (ethyl acetate: *n*-hexane; 8:2). Mp. 260 – 262 °C. ^1^H NMR (400 MHz, DMSO-*d_6_*) δ 9.15 (d, *J* = 10.2 Hz, 1H), 9.13 (s, 1H), 8,97 (d, *J* = 10.2 Hz, 1H), 8.10 (d, *J* = 2.1 Hz, 1H), 7.72 (d, *J* = 15 Hz, 1H), 7.56 (dd, *J* = 15.1, 10.2 Hz, 1H), 7.44 (d, *J* = 10.2 Hz, 1H), 7.28 – 7.24 (m, 5H), 4.75 – 4.73 (m, 1H), 3.11 (dd, *J* =13.8, 5.0 Hz, 1H), 2.87 (dd, *J* = 9.5, 5.0 Hz, 1H), 1.51 – 1.49 (m, 2H), 1.09 – 1.07 (m, 1H). ^13^C NMR (100 MHz, DMSO-*d_6_*) δ 172.34, 166.33, 151.74, 148.36, 141.88, 137.63, 134.53, 129.45, 128.42, 128.00, 127.86, 126.74, 122.89, 120.91, 119.69, 54.56, 37.26, 20.02, 15.89. FT-IR (KBr, cm^−1^) 3254.05, 2926.14, 2247.17, 1672.36, 1668.36, 1523.83, 846.79, 831.36. HRMS (+) Calc. for [C_23_H_20_ClN_4_O_2_]^+^ 418.12748, found: 419.12813 [M+H]^+^. HPLC (protocol C, 50:50 ACN: water): t_R_ (min) = 17.32. Purity: 99.9%.

*(S)-7-Chloro-N-(1-((1-cyanocyclopropyl)amino)-4-methyl-1-oxopentan-2-yl)quinoline-4-carboxamide (**14**)*

Method B. Yield 68%. White solid. *R_f_* = 0.7 (ethyl acetate). Mp. 169 – 170 °C. ^1^H NMR (400 MHz, DMSO-*d_6_*) δ 9.10 (s, 1H), 9.05 (s, 1H), 9.02 (d, *J* = 5.5 Hz, 1H) 8.20 (d, *J* = 11.5 Hz, 1H), 8.15 (d, *J* = 2.5 Hz, 1H), 7.72 (dd, *J* = 11.5, 2.5 Hz, 1H), 7.60 (d, *J* = 5.5 Hz, 1H), 4.48 – 4.46 (m, 1H), 1.70 – 1.63 (m, 3H), 1.52 – 1.48 (m, 2H), 1.19 – 1.14 (m, 2H), 0.92 (2d, *J* = 10.5 Hz, 6H). ^13^C NMR (100 MHz, DMSO-*d_6_*) δ 173.63, 166.83, 152.05, 148.70, 142.06, 134.83, 131.26, 128.38, 123.35, 121.21, 120.20, 51.97, 24.87, 23.38, 21.79, 20.34, 16.22, 16.06. FT-IR (KBr, cm^−1^) 3402.4, 3257.7, 3030.1, 2960.7, 2247.0, 1674.2, 1633.7, 1529.5, 1296.1, 831.31. HRMS (+) Calc. for [C_20_H_22_ClN_4_O_2_]^+^ 384.14313, found: 385.14503 [M+H]^+^. HPLC (protocol C, 65:35 ACN: water): t_R_ (min) = 11.81. Purity: 96.2%.

*(S)-7-Chloro-N-(1-((1-cyanocyclopropyl)amino)-3-(1H-indol-3-yl)-1-oxopropan-2-yl)quinoline-4-carboxamide (**15**)*

Method B. Yield 49%. Yellowish solid. *R_f_* = 0.3 (ethyl acetate). Mp. 196 – 197 °C. ^1^H NMR (400 MHz, DMSO-*d_6_*) δ 10.88 (s, 1H), 9.15 (s, 1H), 9.08 (d, *J* = 10.2 Hz, 1H), 8.97 (d, *J* = 4.2 Hz, 1H), 8.10 (d, *J* = 2.2 Hz, 1H), 7.68 (t, *J* = 7.5 Hz, 1H), 7.54 – 7.52 (m, 1H), 7.46 (d, *J* = 4.5 Hz, 1H), 7.38 (d, *J* = 10.2 Hz 1H), 7.16 (d, *J* = 2.4 Hz, 1H), 7.09 (t, *J* = 5.5 Hz, 1H), 7.00 (t, *J* = 5.8 Hz, 1H), 4.81 – 4.79 (m, 1H), 3.20 (dd, *J* = 13.8, 5.0 Hz, 1H), 3.05 (dd, *J* = 9.5, 5.0 Hz, 1H), 1.51 – 1.48 (m, 2H), 1.11 – 1.08 (m, 2H). ^13^C NMR (100 MHz, DMSO-*d_6_*) δ 173.02, 166.59, 151.94, 148.61, 142.38, 136.57, 134.72, 128.23, 128.05, 128.03, 127.52, 124.56, 123.19, 121.46, 121.26, 119.97, 119.03, 118.76, 111.81, 110.02, 54.26, 27.79, 20.33, 16.20. FT-IR (KBr, cm^−1^) 3254.05, 2926.14, 2247.17, 1672.36, 1665.34, 1522.83, 831.30, 732.98. HRMS (+) Calc. for [C_25_H_21_ClN_5_O_2_]^+^ 458.13838, found: 458.13703 [M+H]^+^. HPLC (protocol C, 50:50 ACN: water): t_R_ (min) = 18.64. Purity: 99.6%.

*(S)-3-((1-((1-Cyanocyclopropyl)amino)-1-oxo-3-phenylpropan-2-yl)carbamoyl)phenyl benzoate (**16**)*

Method B. Yield 89%. White solid. *R_f_* = 0.7 (ethyl acetate: *n*-hexane; 6:4). Mp. 185 – 187 °C. ^1^H NMR (400 MHz, DMSO-*d_6_*) δ 9.05 (s, 1H), 8.79 (d, *J* = 8.0 Hz, 1H), 8.18 – 8.16 (m, 2H), 7.81 – 7.78 (m, 3H), 7.64 (t, *J* = 8.5 Hz, 2H), 7.57 (t, *J* = 8.5 Hz, 2H), 7.50 – 7.48 (m, 1H), 7.31 – 7.23 (m, 4H), 7.18 (tt, *J* = 7.0, 1.5 Hz, 1H), 4.64 – 4.59 (m, 1H), 3.06 (dd, *J* = 13.6, 5.0 Hz, 1H), 3.00 (dd, *J* = 13.5, 8.5 Hz, 1H), 1.49 – 1.45 (m, 2H), 1.06 – 1.01 (m, 2H). ^13^C NMR (100 MHz, DMSO-*d_6_*) δ 173.04, 165.60, 165.02, 150.88, 138.21, 135.72, 134.66, 130.27, 129.98, 129.57, 129.51, 129.16, 128.57, 126.83, 125.69, 125.53, 121.47, 121.15, 55.10, 37.36, 20.18, 16.15, 16.10, 14.55. FT-IR (KBr, cm^−1^) 3254.05, 2926.14, 2247.17, 1672.36, 1668.36, 1523.83, 846.79, 831.36. HRMS (+) Calc. for [C_27_H_24_N_3_O_4_]^+^ 453.17668, found: 454.17800 [M+H]^+^. HPLC (protocol C, 65:35 ACN: water): t_R_ (min) = 19.62. Purity: 99.9%.

*(S)-6-Amino-N-(1-((1-cyanocyclopropyl)amino)-4-methyl-1-oxopentan-2-yl)nicotinamide (**17**)*

Method C. Yield 48%. Yellowish solid. *R_f_* = 0.3 (ethyl acetate: methanol; 8:2). Mp. 100 – 101 °C. ^1^H NMR (200 MHz, CD_3_OD) δ 8.43 (s, 1H), 7.88 (d, *J* = 8.8, 2.1 Hz, 1H), 6.54 (d, *J* = 8.9 Hz,1H), 4.53 – 4.46 (m, 1H), 1.82 – 1.51 (m, 3H), 1.47 – 1.43 (m, 2H), 1.24 – 1.17 (m, 2H), 0.96 – 0.93 (m, 6H). ^13^C NMR (50 MHz, CD_3_OD) δ 176.59, 168.58, 162.86, 149.21, 138.44, 121.42, 119.42, 109.12, 41.60, 26.13 23.54, 21.98, 21.41, 17.15, 16.78. FT-IR (KBr, cm^−1^) 3288.99, 2953.92, 2917.14, 2214.31,1647.63, 1496.06, 1409.33, 1294.89, 1202.33, 1075.33, 1030.81, 922.54, 763.49, 667.21. ESI-MS (+) Calc. for [C_16_H_22_N_5_O_2_]^+^ 316.17735, found: 316.17713 [M+H]^+^. HPLC (protocol B): t_R_ (min) = 8.77. Purity: 97.0%.

*(S)-N-(1-((1-Cyanocyclopropyl)amino)-4-methyl-1-oxopentan-2-yl)-1H-pyrrolo*[*2,3-b*]*pyridine-5-carboxamide (**18**)*

Method C. Yield 45%. White solid. *R_f_* = 0.3 (ethyl acetate: methanol; 8:2). Mp. 79 – 80 °C. ^1^H NMR (200 MHz, CD_3_OD) δ 8.71 (d, *J* = 1.8 Hz, 1H), 8.47 (d, *J* = 2.0 Hz, 1H), 7.46 (d, *J* = 3.5 Hz, 1H), 6.57 (d, *J* = 3.5 Hz, 1H), 4.59 – 4.57 (m, 1H), 1.91 – 1.56 (m, 3H), 1.54 – 1.49 (m, 2H), 1.29 – 1.25 (m, 2H), 1.01 – 0. 89 (m, 6H). ^13^C NMR (50 MHz, CD_3_OD) δ 176.50, 169.61, 163.45, 150.38, 143.32, 129.52, 128.65, 123.21, 120.94, 102.66, 53.50, 38.92, 26.18, 23.41, 22.02, 21.25, 17.17, 16.49. FT-IR (KBr, cm^−1^) 3286.44, 2933.89, 2924.14, 2216.51, 1647.63, 1588.10, 1496.06, 1409.33, 1294.89, 1202.33, 1075.33, 1030.31, 922.54, 763.54, 667.21. ESI-MS (+) Calc. for [C_18_H_22_N_5_O_2_]^+^ 340.17753, found: 340.17689 [M+H]^+^. HPLC (protocol B): t_R_ (min) = 5.38. Purity: 99.0%.

#### Synthesis and characterization of compound 19

Compound **19** has been synthesized from compound **12** by removal of the benzyl protecting group under mild conditions (protocol B).

*3-(tert-Butyl)-N-((2S,3R)-1-((1-cyanocyclopropyl)amino)-3-hydroxy-1-oxobutan-2-yl)-1-methyl-1H-pyrazole-5-carboxamide (19)*

Yield 40%. Yellowish solid. *R_f_* = 0.3 (ethyl acetate). Mp. 201 – 202 °C. ^1^H NMR (200 MHz, CD_3_OD) δ 6.81 (s, 1H), 4.40 (d, *J* = 4.4 Hz, 1H), 4.23 – 4.18 (m, 1H), 4.04 (s, 3H), 1.52 – 1.47(m, 2H), 1.32 (s, 9H), 1.29 – 1.18 (m, 5H). ^13^C NMR (50 MHz, CD_3_OD) δ 174.94, 163.34, 162.74, 137.60, 122.22, 106.16, 69.42, 61.17, 39.91, 34.00, 31.94, 22.41, 21.45, 18.11, 17.83. FT-IR (KBr, cm^−1^) 3288.89, 2953.92, 2917.14, 2214.31, 1646.31, 1589.10, 1495.12, 1049.31, 1294.89, 1200.91, 1033.97, 922.51, 755.51, 681.95. ESI-MS (-) Calc. for [C_17_H_25_N_5_O_3_]^+^ 348.20356, found: 348.20314 [M+H]^+^. HPLC (protocol B): t_R_ (min) = 7.74. Purity: 98.0%.

#### Synthesis of compounds 20-27

Compounds **20***-***27** have been synthesized from the equivalent amino Boc-protected amino acid following the general procedure for amide synthesis (method A).

*(S)-tert-Butyl (1-amino-1-oxo-3-phenylpropan-2-yl)carbamate (**20**)*

Yield 77%. White solid. *R_f_* = 0.6 (ethyl acetate). Mp. 146–149 °C. ^1^H NMR (200 MHz, CD_3_OD) δ 7.26 – 7.16 (m, 5H), 4.44 – 4.41 (m, 1H), 3.14 (dd, *J* = 13.7, 6.0 Hz, 1H), 2.88 (dd, *J* = 13.7, 8.1 Hz, 1H), 1.23 (s, 9H). ^13^C NMR (50 MHz, CD_3_OD) δ 173.98, 155.60, 130.73, 124.93, 122.46, 118.22, 80.20, 54.02, 37.73, 27.94. ESI-MS (+) Calc. for [C_14_H_20_N_2_O_3_] 264.31, found: 287.3 [M+Na]^+^.

*(R)-tert-butyl (1-amino-1-oxo-3-phenylpropan-2-yl)carbamate (**21**)*

Yield 79%. White solid. *R_f_* = 0.6 (ethyl acetate). Mp. 141–142 °C. ^1^H NMR (200 MHz, CD_3_OD) δ 7.24 – 7.17 (m, 5H), 4.42 – 4.37 (m, 1H), 3.14 (dd, *J* = 13.7, 6.0 Hz, 1H), 2.92 (dd, *J* = 13.7, 8.1 Hz, 1H), 1.43 (s, 9H). ^13^C NMR (50 MHz, CD_3_OD) δ 173.12, 154.48, 133.36, 122.46, 121.75, 119.38, 83.51, 55.61, 38.38, 28.66. ESI-MS (+) Calc. for [C_14_H_20_N_2_O_3_] 264.31, found: 287.3 [M+Na]^+^.

*(S)-tert-Butyl (1-amino-4-methyl-1-oxopentan-2-yl)carbamate (**22**)*

Yield 74%. White solid. *R_f_* = 0.4 (ethyl acetate). Mp. 138–141 °C. ^1^H NMR (200 MHz, CDCl_3_) δ 6.58 (s br, 1H), 6.13 (s br, 1H), 5.25 – 4.97 (m, 2H), 4.15 (s br, 1H), 1.73 – 1.43 (m, 3H), 1.41 (s, 9H), 0.92 (d, *J* = 3.2 Hz, 6H). ^13^C NMR (50 MHz, CDCl_3_) δ 172.34, 155.95, 71.56, 28.57, 28.18, 25.08, 23.21, 19.25. ESI-MS (+) Calc. for [C_11_H_22_N_2_O_3_] 230.30, found: 253.3 [M+Na]^+^.

*(R)-tert-Butyl (1-amino-4-methyl-1-oxopentan-2-yl)carbamate (**23**)*

Yield 71%. White solid. *R_f_* = 0.4 (ethyl acetate). Mp. 138–141 °C. ^1^H NMR (200 MHz, CDCl_3_) δ 6.64 (s br, 1H), 6.21 (s br, 1H), 5.21 – 4.98 (m, 2H), 4.16 (s br, 1H), 1.71 – 1.45 (m, 3H), 1.42 (s, 9H), 0.93 (d, *J* = 3.2 Hz, 6H). ^13^C NMR (50 MHz, CDCl_3_) δ 172.11, 155.37, 71.64, 28.61, 28.29, 25.15, 23.34, 19.30. ESI-MS (+) Calc. for [C_11_H_22_N_2_O_3_] 230.30, found: 253.3 [M+Na]^+^.

*(S)-tert-Butyl (1-amino-3-(3-chlorophenyl)-1-oxopropan-2-yl)carbamate (**24**)*

Yield 74%. White solid. *R_f_* = 0.4 (ethyl acetate). Mp. 111–112 °C. ^1^H NMR (200 MHz, CDCl_3_) δ 7.26 – 7.13 (m, 4H), 6.16 (s, 1H), 5.75 (s, 1H), 5.21 (d, *J* = 7.8 Hz, 1H), 4.42 – 4.36 (m, 1H), 3.19 – 2.94 (m, 2H), 1.44 (s, 9H). ^13^C NMR (50 MHz, CDCl_3_) δ 173.53, 155.16, 138.90, 134.09, 130.04, 129.44, 127.55, 127.11, 80.31, 37.88, 28.16. ESI-MS (+) Calc. for [C_14_H_19_ClN_2_O_3_] 298.77, found: 321.8 [M+Na]^+^.

*(S)-tert-Butyl (1-amino-1-oxo-3-(pyridin-4-yl)propan-2-yl)carbamate (**25**)*

Yield 92%. White solid. *R_f_* = 0.2 (ethyl acetate). Mp. 131–133 °C. ^1^H NMR (200 MHz, CDCl_3_) δ 8.42 (d, *J* = 5.7 Hz, 2H), 7.22 (d, *J* = 5.9 Hz, 2H), 4.35 – 4.31 (m, 1H), 3.18 (dd, *J* = 13.7, 6.0 Hz, 1H), 2.81 (dd, *J* = 13.7, 8.1 Hz, 1H), 1.49–1.33 (s br, 9H). ^13^C NMR (50 MHz, CDCl_3_) δ 173.69, 155.86, 148.48, 147.61, 118.42, 80.22, 54.51, 37.77, 27.89. ESI-MS (+) Calc. for [C_13_H_19_N_3_O_3_] 265.31, found: 266.2 [M+H]^+^.

*tert-Butyl ((2S,3R)-1-amino-3-(benzyloxy)-1-oxobutan-2-yl)carbamate (**26**)*

Yield 65%. White solid. *R_f_* = 0.5 (ethyl acetate). Mp. 145–147 °C. ^1^H NMR (500 MHz, CDCl_3_) δ 7.37 – 7.28 (m, 5H), 6.50 (s, 1H), 5.70 – 5.51 (m, 2H), 4.62 (q, *J* = 4.6 Hz, 2H), 4.34 – 4.31 (m, 1H), 4.18 – 4.11 (m, 1H), 1.45 (s, 9H), 1.19 (d, *J* = 6.3 Hz, 3H). ^13^C NMR (125 MHz, CDCl_3_) δ 172.09, 155.66, 137.78, 128.41, 127.82, 127.73, 80.03, 74.53, 71.61, 57.13, 28.23, 27.84, 18.92. ESI-MS (+) Calc. for [C_16_H_24_N_2_O_4_] 308.37, found: 331.4 [M+Na]^+^.

*tert-Butyl ((2R,3S)-1-amino-3-(benzyloxy)-1-oxobutan-2-yl)carbamate (**27**)*

Yield 68%. White solid. *R_f_* = 0.5 (ethyl acetate). Mp. 143–145 °C. ^1^H NMR (500 MHz, CDCl_3_) δ 7.37 – 7.28 (m, 5H), 6.50 (s, 1H), 5.70 – 5.51 (m, 2H), 4.62 (q, *J* = 4.6 Hz, 2H), 4.33 (s br, 1H), 4.27 – 4.19 (m, 1H), 1.45 (s, 9H), 1.19 (d, *J* = 6.3 Hz, 3H). ^13^C NMR (125 MHz, CDCl_3_) δ 172.74, 156.31, 138.44, 129.06, 128.47, 128.38, 80.68, 77.16, 75.18, 72.26, 57.78, 28.89, 28.49, 19.57. ESI-MS (+) Calc. for [C_16_H_24_N_2_O_4_] 308.37, found: 331.4 [M+Na]^+^.

#### Synthesis of compounds 28-38

Compounds **28**-**38** have been synthesized in two steps from their precursors **20**-**27**. After removal of the Boc-protecting group (method B), the resulting free amine was coupled to the carboxylic acid following the general procedure for amide synthesis (method B).

*tert-Butyl ((S)-1-(((S)-1-amino-1-oxo-3-phenylpropan-2-yl)amino)-1-oxo-3-phenylpropan-2-yl)carbamate (**28**)*

Yield 92%. White solid. *R_f_* = 0.8 (ethyl acetate). Mp. 186–188 °C. ^1^H NMR (200 MHz, CD_3_OD) δ 7.37 – 7.22 (m, 6H), 7.16 – 7.09 (m, 4H), 4.76 – 4.63 (m, 1H), 4.34 – 4.20 (m, 1H), 3.15 – 2.95 (m, 2H), 2.91 – 2.62 (m, 2H), 1.39 (s, 9H). ^13^C NMR (50 MHz, CD_3_OD) δ 174.36, 172.35, 156.09, 136.44, 129.04, 128.35, 128.27, 126.78, 126.64, 79.89, 56.00, 53.78, 37.73, 37.24, 27.84. ESI-MS (+) Calc. for [C_23_H_29_N_3_O_4_] 411.49, found: 434.3 [M+Na]^+^.

*tert-Butyl ((S)-1-(((R)-1-amino-1-oxo-3-phenylpropan-2-yl)amino)-1-oxo-3-phenylpropan-2-yl)carbamate (**29**)*

Yield 92%. White solid. *R_f_* = 0.8 (ethyl acetate). Mp. 185–187 °C. ^1^H NMR (200 MHz, CD_3_OD) δ 7.75 – 7.60 (m, 6H), 7.56 – 7.49 (m, 4H), 5.14 – 5.01 (m, 1H), 4.72 – 4.57 (m, 1H), 3.53 – 3.33 (m, 2H), 3.29 – 3.00 (m, 2H), 1.77 (s, 9H). ^13^C NMR (50 MHz, CD_3_OD) δ 174.85, 172.83, 156.58, 136.93, 129.53, 128.83, 128.76, 127.26, 127.13, 80.38, 56.49, 54.26, 38.22, 37.73, 28.33. ESI-MS (+) Calc. for [C_23_H_29_N_3_O_4_] 411.49, found: 434.3 [M+Na]^+^.

*tert-Butyl ((S)-1-(((S)-1-amino-4-methyl-1-oxopentan-2-yl)amino)-1-oxo-3-phenylpropan-2-yl)carbamate (**30**)*

Yield 83%. White solid. *R_f_* = 0.8 (ethyl acetate). Mp. 164–165 °C. ^1^H NMR (200 MHz, CD_3_OD) δ 7.24 – 7.20 (m, 5H), 4.33 – 4.28 (m, 1H), 4.11 – 4.05 (m, 1H), 3.03 – 2.93 (m, 2H), 1.56 – 1.38 (m, 2H), 1.36 (s, 9H), 1.26 – 1.24 (m, 1H), 0.89 (d, *J* = 2.6 Hz, 6H). ^13^C NMR (50 MHz, CD_3_OD) δ 175.39, 154.59, 143.42, 136.55, 128.99, 128.29, 126.65, 84.91, 72.86, 52.36, 51.47, 27.61, 24.41, 22.20, 21.01. ESI-MS (+) Calc. for [C_20_H_31_N_3_O_4_] 377.48, found: 400.5 [M+Na]^+^.

*tert-Butyl ((S)-1-(((R)-1-amino-4-methyl-1-oxopentan-2-yl)amino)-1-oxo-3-phenylpropan-2-yl)carbamate (**31**)*

Yield 88%. White solid. *R_f_* = 0.8 (ethyl acetate). Mp. 168–170 °C. ^1^H NMR (200 MHz, CD_3_OD) δ 7.24 – 7.21 (m, 5H), 4.31 – 4.21 (m, 1H), 4.11 – 4.09 (m, 1H), 3.04 – 2.94 (m, 2H), 1.57 – 1.40 (m, 2H), 1.39 (s, 9H), 1.27 – 1.24 (m, 1H), 0.90 (d, *J* = 2.6 Hz, 6H). ^13^C NMR (50 MHz, CD_3_OD) δ 175.80, 155.00, 143.83, 136.96, 129.40, 128.70, 127.06, 85.32, 77.16, 73.27, 52.78, 51.89, 28.02, 24.82, 22.61, 21.42. ESI-MS (+) Calc. for [C_20_H_31_N_3_O_4_] 377.48, found: 400.5 [M+Na]^+^.

*tert-Butyl ((S)-1-(((S)-1-amino-3-(3-chlorophenyl)-1-oxopropan-2-yl)amino)-1-oxo-3-phenylpropan-2-yl)carbamate (**32**)*

Yield 78%. White solid. *R_f_* = 0.6 (ethyl acetate). Mp. 148–150 °C. ^1^H NMR (200 MHz, CDCl_3_) δ 7.38 – 7.26 (m, 7H), 7.10 – 7.06 (m, 2H), 6.63 (s br, 1H), 6.39 (s br, 1H), 4.81 – 4.77 (m, 1H), 4.38 – 4.35 (m, 1H), 3.05 – 2.97 (m, 4H), 1.41 (s, 9H). ^13^C NMR (50 MHz, CDCl_3_) δ 172.83, 161.82, 138.54, 136.14, 134.46, 133.26, 130.05, 129.51, 129.36, 129.02, 127.69, 127.36, 126.78, 80.95, 62.51, 53.21, 52.76, 38.74, 28.23. ESI-MS (+) Calc. for [C_23_H_28_ClN_3_O_4_] 445.94, found: 468.8 [M+Na]^+^.

*tert-Butyl ((S)-1-(((S)-1-amino-1-oxo-3-(pyridin-4-yl)propan-2-yl)amino)-1-oxo-3-phenylpropan-2-yl)carbamate (**33**)*

Yield 71%. Yellowish solid. *R_f_* = 0.4 (ethyl acetate). Mp. 155–157 °C. ^1^H NMR (200 MHz, CD_3_OD) δ 8.63 (d, *J* = 5.7 Hz, 2H), 7.58 – 7.30 (m, 7H), 4.94 – 4.89 (1H, m), 4.55 – 4.40 (m, 1H), 3.44 (dd, *J* = 13.8, 5.5 Hz, 2H), 3.28 – 2.93 (m, 2H), 1.60 (s, 9H). ^13^C NMR (50 MHz, CDCl_3_) δ 173.27, 172.38, 148.58, 140.95, 136.44, 135.10, 128.97, 128.32, 126.68, 124.97, 99.74, 55.40, 27.73, 23.92, 13.06, 10.47.ESI-MS (+) Calc. for [C_23_H_28_ClN_3_O_4_] 445.94, found: 468.8 [M+Na]^+^.

*tert-Butyl ((S)-1-(((2S,3R)-1-amino-3-(benzyloxy)-1-oxobutan-2-yl)amino)-1-oxo-3-phenylpropan-2-yl)carbamate (**34**)*

Yield 88%. Yellowish solid. *R_f_* = 0.8 (ethyl acetate). Mp. 161–163 °C. ^1^H NMR (200 MHz, CDCl_3_) δ 7.38 – 7.26 (m, 7H), 7.10 – 7.06 (m, 2H), 6.63 (s br, 1H), 6.39 (s br, 1H), 5.65 (s br, 1H), 5.08 (s br, 1H), 4.81 – 4.77 (m, 1H), 4.38 – 4.35 (m, 1H), 3.15 – 2.97 (m, 5H), 1.74 – 1.47 (m, 3H), 1.41 (s, 9H). ^13^C NMR (50 MHz, CDCl_3_) δ 172.98, 171.90, 156.38, 137.70, 136.53, 128.99, 128.42, 128.13, 127.64, 127.58, 126.75, 80.30, 74.03, 71.36, 60.44, 38.05, 27.67, 20.33, 15.75. ESI-MS (+) Calc. for [C_25_H_33_N_3_O_5_] 455.55, found: 478.5 [M+Na]^+^.

*tert-Butyl ((S)-1-(((2R,3S)-1-amino-3-(benzyloxy)-1-oxobutan-2-yl)amino)-1-oxo-3-phenylpropan-2-yl)carbamate (**35**)*

Yield 78%. Yellowish solid. *R_f_* = 0.8 (ethyl acetate). Mp. 138–139 °C. ^1^H NMR (200 MHz, CDCl_3_) δ 7.40 – 7.37 (m, 7H), 7.11 – 7.08 (m, 2H), 6.58 (s br, 1H), 6.43 (s br, 1H), 5.61 (s br, 1H), 5.23 (s br, 1H), 4.83 – 4.78 (m, 1H), 4.34 – 4.31 (m, 1H), 3.32 – 2.85 (m, 5H), 1.39 (s, 9H), 1.19 – 1.09 m, 3H). ^13^C NMR (50 MHz, CDCl_3_) δ 173.34, 172.27, 156.74, 138.07, 136.90, 129.35, 128.79, 128.49, 128.01, 127.95, 127.12, 80.66, 74.40, 71.73, 60.81, 38.42, 28.04, 20.70, 16.12.ESI-MS (+) Calc. for [C_25_H_33_N_3_O_5_] 455.55, found: 478.5 [M+Na]^+^.

*tert-Butyl ((S)-1-(((2S,3R)-1-amino-3-(benzyloxy)-1-oxobutan-2-yl)amino)-4-methyl-1-oxopentan-2-yl)carbamate (**36**)*

Yield 76%. Yellowish solid. *R_f_* = 0.8 (ethyl acetate). Mp. 88–91 °C. ^1^H NMR (200 MHz, CDCl_3_) δ 7.30 – 7.22 (m, 5H), 7.19 – 7.15 (m, 1H), 6.24 (s br, 1H), 6.04 (s br, 1H), 5.36 (s br, 2H), 4.57 (s, 2H), 4.56 – 4.52 (m, 1H), 4.36 – 4.21 (m, 1H), 4.17 – 4.11 (m, 1H), 1.77 – 1.48 (m, 3H), 1.38 (s, 9H), 1.13 (d, *J* = 6.4 Hz, 3H), 0.94 – 0.90 (m, 6H). ^13^C NMR (50 MHz, CDCl_3_) δ 172.84, 171.86, 155.98, 137.88, 128.29, 127.66, 127.63, 80.38, 73.89, 71.58, 56.40, 54.01, 40.85, 28.12, 24.74, 22.92, 21.62. ESI-MS (+) Calc. for [C_22_H_35_N_3_O_5_] 421.53, found: 444.7 [M+Na]^+^.

*tert-Butyl ((S)-1-(((2R,3S)-1-amino-3-(benzyloxy)-1-oxobutan-2-yl)amino)-4-methyl-1-oxopentan-2-yl)carbamate (**37**)*

Yield 73%. Yellowish solid. *R_f_* = 0.8 (ethyl acetate). Mp. 86–88 °C. ^1^H NMR (200 MHz, CDCl_3_) δ 7.31 – 7.22 (m, 5H), 7.20 – 7.11 (m, 1H), 6.26 (s br, 1H), 6.02 (s br, 1H), 5.36 (s br, 2H), 4.54 (s, 2H), 4.56 – 4.53 (m, 1H), 4.33 – 4.28 (m, 1H), 4.16 – 4.11 (m, 1H), 1.76 – 1.48 (m, 3H), 1.48 (s, 9H), 1.11 (d, *J* = 6.4 Hz, 3H), 0.96 – 0.90 (m, 6H). ^13^C NMR (50 MHz, CDCl_3_) δ 172.84, 171.86, 155.98, 137.88, 128.29, 127.66, 127.63, 80.38, 73.89, 71.58, 56.40, 54.01, 40.85, 28.12, 24.74, 22.92, 21.62. ESI-MS (+) Calc. for [C_22_H_35_N_3_O_5_] 421.53, found: 444.5 [M+Na]^+^.

*tert-Butyl ((S)-1-(((2S,3R)-1-amino-3-(benzyloxy)-1-oxobutan-2-yl)amino)-3-(3-chlorophenyl)-1-oxopropan-2-yl)carbamate (**38**)*

Yield 77%. Yellowish solid. *R_f_* = 0.6 (ethyl acetate). Mp. 164–165 °C. ^1^H NMR (200 MHz, CD_3_OD) δ 7.47 – 7.22 (m, 9H), 4.55 – 4.37 (m, 1H), 4.29 – 4.17 (m, 2H), 3.26 (dd, *J* = 13.9, 5.1 Hz, 1H), 3.07 – 2.96 (m, 1H), 2.92 (s, 2H), 1.44 (s, 9H), 1.27 (d, *J* = 6.3 Hz, 3H). ^13^C NMR (50 MHz, CD_3_OD) δ 172.84, 171.76, 156.24, 138.90, 137.56, 136.39,130.04, 128.85, 128.28, 127.99, 127.50, 127.44, 126.61, 80.16, 73.89, 71.22, 60.30, 49.00, 37.92, 27.53, 20.19. ESI-MS (+) Calc. for [C_25_H_32_ClN_3_O_5_] 489.99, found: 513.1 [M+Na]^+^.

#### Synthesis of compounds 39-49

Compounds **39**-**49** have been synthesized in two steps from their precursors **28**-**38**. After removal of the Boc-protecting group (method B), the free amine was coupled to the carboxylic acid following the general procedure for amide synthesis (method C).

*N-((S)-1-(((S)-1-Amino-1-oxo-3-phenylpropan-2-yl)amino)-1-oxo-3-phenylpropan-2-yl)-3-(tert-butyl)-1-methyl-1H-pyrazole-5-carboxamide (**39**)*

Yield 79%. Yellowish solid. *R_f_* = 0.3 (ethyl acetate). Mp. 109–110 °C. ^1^H NMR (200 MHz, CDCl_3_) δ7.32 – 6.96 (m, 10H), .6.65 (s br, 1H), 6.26 (s, 1H), 6.09 (s br, 1H), 4.68 – 4.58 (m, 2H), 3.81 (s, 3H), 2.87 – 2.74 (m, 4H), 1.11 (s, 9H). ^13^C NMR (50 MHz, CDCl_3_) δ 173.37, 165.56, 162.46, 160.20, 159.99, 136.34, 136.16, 134.40, 129.13, 128.39, 126.87, 126.81, 124.46, 103.30, 54.93, 53.91, 38.44, 36.34, 31.69, 31.28, 30.27. ESI-MS (+) Calc. for [C_27_H_33_N_5_O_3_] 475.58, found: 498.7 [M+Na]^+^.

*N-((S)-1-(((R)-1-Amino-1-oxo-3-phenylpropan-2-yl)amino)-1-oxo-3-phenylpropan-2-yl)-3-(tert-butyl)-1-methyl-1H-pyrazole-5-carboxamide (**40**)*

Yield 72%. Yellowish solid. *R_f_* = 0.4 (ethyl acetate). Mp. 121–122 °C. ^1^H NMR (200 MHz, CDCl_3_) δ 7.59 – 7.13 (m, 10H), 6.88 (s, 1H), 6.50 (s, 1H), 6.32 (s br, 1H), 4.92 – 4.82 (m, 2H), 4.04 (s, 3H), 3.10 – 2.91 (m, 4H), 1.34 (s, 9H). ^13^C NMR (50 MHz, CDCl_3_) δ 172.94, 165.13, 162.04, 159.77, 159.56, 135.92, 135.74, 133.97, 128.70, 127.97, 126.45, 126.38, 124.04, 102.88, 54.50, 53.48, 38.01, 35.91, 31.26, 30.86, 29.84. ESI-MS (+) Calc. for [C_27_H_33_N_5_O_3_] 475.58, found: 498.7 [M+Na]^+^.

*N-((S)-1-(((S)-1-Amino-4-methyl-1-oxopentan-2-yl)amino)-1-oxo-3-phenylpropan-2-yl)-3-(tert-butyl)-1-methyl-1H-pyrazole-5-carboxamide (**41**)*

Yield 85%. Yellowish solid. *R_f_* = 0.3 (ethyl acetate). Mp. 152–153 °C. ^1^H-NMR (200 MHz, CDCl_3_) δ. 7.42 – 7.25 (m, 5H), 6.81 (s, 1H), 4.77 – 4.65 (m, 1H), 4.36 – 4.08 (m, 1H), 4.08 (s, 3H), 3.22 – 3.18 (m, 2H), 1.89 – 1.46 (m, 3H), 1.40 (s, 9H), 0.88 – 0.80 (m, 6H). ^13^C NMR (50 MHz, CDCl_3_) δ 173.57, 168.11, 162.00, 161.25, 137.23, 135.94, 130.02, 129.31, 127.64, 104.97, 57.07, 49.00, 38.87, 32.59, 31.75, 30.78, 24.80, 23.57, 21.42. ESI-MS (+) Calc. for [C_24_H_35_N_5_O_3_] 441.57, found: 464.5 [M+Na]^+^.

*N-((S)-1-(((S)-1-Amino-3-(3-chlorophenyl)-1-oxopropan-2-yl)amino)-1-oxo-3-phenylpropan-2-yl)-3-(tert-butyl)-1-methyl-1H-pyrazole-5-carboxamide (**42**)*

Yield 72%. Yellowish solid. *R_f_* = 0.3 (ethyl acetate). Mp. 131–133 °C. ^1^H NMR (200 MHz, CDCl_3_) δ 7.45 – 7.28 (s, 5H), 6.83 (s, 1H), 4.87 – 4.68 (m, 1H), 4.39 – 4.11 (m, 1H), 4.11 (s, 3H), 3.25 – 3.21 (m, 2H), 1.76 – 1.49 (m, 3H), 1.42 (s, 9H), 0.88 – 0.83 (m, 6H). ^13^C NMR (50 MHz, CDCl_3_) δ 176.49, 171.02, 164.92, 164.17, 140.15, 138.86, 132.94, 132.23, 130.56, 107.89, 76.35, 59.99, 41.79, 35.51, 34.67, 33.70, 27.72, 26.49, 24.34. ESI-MS (+) Calc. for [C_24_H_35_N_5_O_3_] 441.57, found: 464.5 [M+Na]^+^.

*tert-Butyl ((S)-1-(((S)-1-amino-3-(3-chlorophenyl)-1-oxopropan-2-yl)amino)-1-oxo-3-phenylpropan-2-yl)carbamate (**43**)*

Yield 80%. Yellowish solid. *R_f_* = 0.2 (ethyl acetate). Mp. 101–103 °C. ^1^H NMR (200 MHz, CDCl_3_:CD_3_OD 10:1) δ. 7.58 – 6.99 (m, 9H), 6.47 (s, 1H), 4.77 – 4.62 (m, 2H), 4.04 (s, 3H), 3.39 – 2.91 (m, 4H), 1.34 (s, 9H). ^13^C NMR (50 MHz, CDCl_3_:CD_3_OD 10:1) δ 172.23, 172.21, 161.25, 161.03, 137.40, 136.96, 135.83, 135.15, 130.68, 129.87, 129.23, 128.42, 128.30, 127.69, 117.62, 104.55, 55.12, 49.00, 42.41, 38.76, 38.62, 32.51, 30.78. ESI-MS (+) Calc. for [C_27_H_32_ClN_5_O_3_] 510.03, found: 532.9 [M+Na]^+^.

*N-((S)-1-(((S)-1-Amino-1-oxo-3-(pyridin-4-yl)propan-2-yl)amino)-1-oxo-3-phenylpropan-2-yl)-3-(tert-butyl)-1-methyl-1H-pyrazole-5-carboxamide (**44**)*

Yield 66%. Yellowish oil. *R_f_* = 0.2 (ethyl acetate). ^1^H NMR (200 MHz, CDCl_3_) δ 8.63 (d, *J* = 5.7 Hz, 2H), 7.58 – 7.30 (m, 7H), 7.36 – 7.13 (m, 7H), 6.59 (s, 1H), 4.84 – 4.68 (m, 2H), 3.89 (s, 3H), 3.35 – 3.07 (m, 2H), 1.27 (s, 9H). ^13^C NMR (50 MHz, CDCl_3_) δ 174.42, 173.04, 161.51, 161.12, 149.40, 148.81, 138.04, 136.11, 129.86, 129.09, 127.45, 126.03, 104.55, 55.73, 54.08, 38.35, 37.93, 37.75, 32.45, 30.45. ESI-MS (+) Calc. for [C_26_H_32_N_6_O_3_] 476.57, found:477.3 [M+H]^+^.

*N-((S)-1-(((2S,3R)-1-Amino-3-(benzyloxy)-1-oxobutan-2-yl)amino)-1-oxo-3-phenylpropan-2-yl)-3-(tert-butyl)-1-methyl-1H-pyrazole-5-carboxamide (**45**)*

Yield 65%. Yellowish solid. *R_f_* = 0.3 (ethyl acetate). Mp 109–110 °C. ^1^H NMR (200 MHz, CDCl_3_) δ 7.30 – 7.08 (m, 9H), 6.40 (d, *J* = 14.4 Hz, 1H), 6.26 (s, 1H), 5.00 – 4.76 (m, 1H), 4.52 (s, 2H), 4.17 – 4.06 (m, 2H), 3.98 (s, 3H), 3.20 – 3.09 (m, 2H), 1.27 (s, 9H), 1.11 (d, *J* = 6.2 Hz, 3H). ^13^C NMR (50 MHz, CDCl_3_) δ 171.79, 171.19, 160.58, 160.20, 137.80, 136.25, 134.62, 129.26, 128.78, 128.44, 127.84, 127.78, 127.21, 103.32, 74.00, 71.58, 60.39, 56.64, 38.86, 38.61, 31.91, 30.49, 21.02. ESI-MS (+) Calc. for [C_29_H_37_N_5_O_4_] 519.64, found:549.4 [M+Na]^+^.

*N-((S)-1-(((2R,3S)-1-Amino-3-(benzyloxy)-1-oxobutan-2-yl)amino)-1-oxo-3-phenylpropan-2-yl)-3-(tert-butyl)-1-methyl-1H-pyrazole-5-carboxamide (**46**)*

Yield 61%. Yellowish solid. *R_f_* = 0.3 (ethyl acetate). Mp 119–120 °C. ^1^H NMR (200 MHz, CDCl_3_) δ.7.27 – 7.04 (m, 9H), 6.37 (d, *J* = 14.4 Hz, 1H), 6.22 (s, 1H), 4.97 – 4.73 (m, 1H), 4.48 (s, 2H), 4.14 – 4.03 (m, 2H), 3.95 (s, 3H), 3.17 – 3.06 (dd, *J* = 11.3, 5.1 Hz, 2H), 1.23 (s, 9H), 1.08 (d, *J* = 6.2 Hz, 3H). ^13^C NMR (50 MHz, CDCl_3_) δ 171.79, 171.19, 160.58, 160.20, 137.80, 136.25, 134.62, 129.26, 128.78, 128.44, 127.84, 127.78, 127.21, 103.32, 74.00, 71.58, 60.39, 56.64, 38.86, 38.61, 31.91, 30.49, 21.02. ESI-MS (+) Calc. for [C_29_H_37_N_5_O_4_] 519.64, found: 549.4 [M+Na]^+^.

*N-((S)-1-(((2S,3R)-1-Amino-3-(benzyloxy)-1-oxobutan-2-yl)amino)-4-methyl-1-oxopentan-2-yl)-3-(tert-butyl)-1-methyl-1H3-pyrazole-5-carboxamide (**47**)*

Yield 73%. Yellowish solid. *R_f_* = 0.3 (ethyl acetate). Mp 177–178 °C. ^1^H NMR (200 MHz, CDCl_3_) δ. 7.27 – 7.20 (m, 4H), 7.14 (d, *J* = 8.5 Hz, 1H), 6.42 (s, 1H), 4.85 – 4.82 (m, 1H), 4.62 – 4.52 (s, 2H), 3.96 (s, 3H), 3.87 – 3.84 (m, 1H), 1.69 – 1.59 (m, 3H), 1.22 (s, 9H). 1.18 (d, *J* = 6.3 Hz, 3H), 0.90 (d, *J* = 6.0 Hz, 3H), 0.97 (d, *J* = 6.0 Hz, 3H). ^13^C NMR (50 MHz, CDCl_3_) δ 172.35, 171.66, 160.37, 160.31, 137.82, 134.69, 128.57, 128.00, 127.74, 103.21, 74.07, 71.71, 56.24, 52.12, 39.03, 38.69, 32.02, 30.59, 25.00, 23.01, 22.0. ESI-MS (+) Calc. for [C_26_H_39_N_5_O_4_] 485.62, found: 508.5 [M+Na]^+^.

*N-((S)-1-(((2R,3S)-1-Amino-3-(benzyloxy)-1-oxobutan-2-yl)amino)-4-methyl-1-oxopentan-2-yl)-3-(tert-butyl)-1-methyl-1H3-pyrazole-5-carboxamide (**48**)*

Yield 70%. Yellowish solid. *R_f_* = 0.3 (ethyl acetate). Mp 170–171 °C. ^1^H NMR (200 MHz, CDCl_3_) δ 7.33 – 7.21 (m, 4H), 7.15 (d, *J* = 8.5 Hz, 1H), 6.34 (s, 1H), 4.85-4.82 (m, 1H), 4.64 – 4.53 (s, 2H), 3.98 (s, 3H), 3.88 – 3.86 (m, 1H), 1.71 – 1.60 (m, 3H), 1.24 (s, 9H), 1.21 (d, *J* = 6.3 Hz, 3H), 0.91 (d, *J* = 6.0 Hz, 3H), 0.89 (d, *J* = 6.0 Hz, 3H). ^13^C NMR (50 MHz, CDCl_3_) δ 171.72, 171.02, 159.73, 159.68, 137.18, 134.05, 127.94, 127.36, 127.10, 102.58, 73.43, 71.08, 55.60, 51.49, 38.39, 38.06, 31.39, 29.95, 24.37, 22.38, 21.44. ESI-MS (+) Calc. for [C_26_H_39_N_5_O_4_] 485.62, found: 508.5 [M+Na]^+^.

*N-((S)-1-(((2S,3R)-1-Amino-3-(benzyloxy)-1-oxobutan-2-yl)amino)-3-(3-chlorophenyl)-1-oxopropan-2-yl)-3-(tert-butyl)-1-methyl-1H-pyrazole-5-carboxamide (**49**)*

Yield 65%. Yellowish solid. *R_f_* = 0.2 (ethyl acetate). Mp 102–104 °C. ^1^H NMR (200 MHz, CD_3_OD) δ 8.59 (d, *J* = 7.8 Hz, 1H), 8.03 (d, *J* = 8.4 Hz, 1H), 7.26 (dd, *J* = 5.6, 2.2 Hz, 9H), 6.59 (s, 1H), 4.64 – 4.45 (m, 3H), 4.16 – 4.11 (m, 1H), 3.86 (s, 3H), 3.27 – 3.25 (m, 1H), 3.03 (dd, *J* = 14.7, 10.8 Hz, 1H), 1.28 (s, 9H), 1.22 (d, *J* = 6.3 Hz, 3H). ^13^C NMR (50 MHz, CD_3_OD) δ 174.44, 173.31, 162.05, 161.35, 140.90, 139.31, 136.28, 134.96, 130.78, 130.33, 129.08, 128.62, 128.52, 128.44, 127.71, 104.84, 75.78, 72.24, 58.45, 55.77, 38.46, 37.34, 32.66, 30.66, 16.58. ESI-MS (+) Calc. for [C_29_H_36_ClN_5_O_4_] 554.08, found: 577.1 [M+Na]^+^.

#### Synthesis of compounds 50-60

Compounds **50**-**60** have been synthesized by dehydration of the corresponding primary amide precursor **39**-**49** with cyanuric chloride (method A).

*3-(tert-Butyl)-N-((S)-1-(((S)-1-cyano-2-phenylethyl)amino)-1-oxo-3-phenylpropan-2-yl)-1-methyl-1H-pyrazole-5-carboxamide (**50**)*

Yield 89%. Yellowish solid. *R_f_* = 0.6 (ethyl acetate: *n*-hexane; 6:4). Mp. 89 – 90 °C. ^1^H NMR (400 MHz, CDCl_3_) δ 7.32 – 7.21 (m, 8H), 7.16 (dd, *J* = 6.4, 2.8 Hz, 2H), 6.90 (d, *J* = 8.2 Hz, 1H), 6.82 (d, *J* = 8.2 Hz, 1H), 6.35 (s, 1H), 5.04 – 5.02 (m, 1H), 4.82 – 4.80 (m, 1H), 4.02 (s, 3H), 3.21 – 3.07 (m, 2H), 3.00 – 2.98 (m, 2H), 1.33 – 1.30 (m, 9H). ^13^C NMR (100 MHz, CDCl_3_) δ 170.30, 160.25, 160.30, 135.55, 134.04, 133.37, 129.08, 129.04, 128.75, 128.72, 127.71, 127.26, 117.25, 103.06, 54.17, 41.48, 38.77, 38.35, 37.95, 31.75, 30.26. FT-IR (KBr, cm^−1^) 3300.81, 2951.21, 2909.60, 2243.70, 1677.69, 1648.55, 1544.51, 1278.15, 1232.37, 986.82, 751.92, 670.52, 524.68, 424.97. HRMS (+) Calc. for [C_27_H_32_N_5_O_2_]^+^ 458.25560, found: 458.2586 [M+H]^+^. HPLC (protocol A): t_R_ (min) = 9.07. Purity 99.0%.

*3-(tert-Butyl)-N-((S)-1-(((R)-1-cyano-2-phenylethyl)amino)-1-oxo-3-phenylpropan-2-yl)-1-methyl-1H-pyrazole-5-carboxamide (**51**)*

Yield 92%. Yellowish solid. *R_f_* = 0.6 (ethyl acetate: *n*-hexane; 6:4). Mp. 98 – 99 °C. ^1^H NMR (200 MHz, CDCl_3_) δ 7.34 – 7.12 (m, 9H), 6.91 (d, *J* = 7.9 Hz, 1H), 6.34 (s, 1H), 5.09 – 5.05 (m, 1H), 4.94 – 4.91 (m, 1H), 4.00 (s, 3H), 3.12 (dd, *J* = 13.8, 6.4 Hz, 2H), 2.98 (dd, *J* = 13.8, 6.4 Hz, 2H), 1.30 (s, 9H). ^13^C NMR (50 MHz, CDCl_3_) δ 170.75, 160.43, 160.33, 135.98, 134.32, 133.78, 129.42, 129.33, 129.01, 128.86, 127.99, 127.42, 118.87, 103.48, 54.38, 41.82, 38.96, 38.48, 38.26, 31.98, 30.51. FT-IR (KBr, cm^−1^) 3350.28, 3256.30, 2953.92, 2165.27, 1638.14, 1511.6, 1307.15, 1160.05, 1086.49, 906.70, 820.89, 694.21, 669.69, 543.02. HRMS (+) Calc. for [C_27_H_32_N_5_O_2_]^+^ 458.25560, found: 458.2586 [M+H]^+^. HPLC (protocol A): t_R_ (min) = 17.32. Purity 99.3%.

*3-(tert-Butyl)-N-((S)-1-(((S)-1-cyano-3-methylbutyl)amino)-1-oxo-3-phenylpropan-2-yl)-1-methyl-1H-pyrazole-5-carboxamide (**52**)*

Yield 84%. Yellowish solid. *R_f_* = 0.7 (ethyl acetate: *n*-hexane; 6:4). Mp 81 – 82 °C. ^1^H NMR (200 MHz, CD_3_OD) δ 7.29 – 7.20 (m, 5H), 6.64 (s, 1H), 4.80 – 4.69 (m, 2H), 3.90 (s, 3H), 3.23 – 2.94 (m, 2H), 1.78 – 1.62 (m, 3H), 1.27 (s, 9H), 0.95 – 0.91 (m, 6H). ^13^C NMR (50 MHz, CD_3_OD) δ 173.33, 161.87, 161.51, 138.07, 136.59, 130.36, 129.56, 127.93, 119.68, 105.02, 56.04, 49.00, 42.00, 40.04, 38.69, 32.86, 30.88, 25.80, 22.35, 22.19. FT-IR (KBr, cm^−1^) 3403.41, 3309.42, 3203.18, 2953.92, 2937.57, 1695.35, 1634.05, 1523.72, 1368.44, 1274.46, 1245.86, 1168.22, 835.58, 702.38, 669.69, 567.54. HRMS (+) Calc. for [C_24_H_34_N_5_O_2_]^+^ 423.27125, found: 424.27572 [M+H]^+^. HPLC (protocol A): t_R_ (min)= 9.16. Purity 97.4%.

*3-(tert-Butyl)-N-((S)-1-(((S)-1-cyano-3-methylbutyl)amino)-1-oxo-3-phenylpropan-2-yl)-1-methyl-1H-pyrazole-5-carboxamide (**53**)*

Yield 79%. White solid. *R_f_* = 0.7 (ethyl Acetate: *n*-hexane; 6:4). Mp. 91 – 94 °C. ^1^H NMR (200 MHz, CDCl_3_) δ 7.32 – 7.21 (m, 3H), 6.98 (d, *J* = 8.2 Hz, 1H), 6.88 (d, *J* = 8.2 Hz, 1H), 6.34 (s, 1H), 4.92 – 4.86 (m, 1H), 4.77 – 4.71 (m, 1H), 3.99 (s, 3H), 3.29 – 3.19 (m, 2H), 1.63 – 1.46 (m, 3H), 1.27 (s, 9H), 0.88 (2d, *J* = 5.9 Hz, 6H). ^13^C NMR (50 MHz, CDCl_3_) δ 171.28, 161.15, 161.04, 136.68, 135.04, 129.97, 129.56, 128.05, 119.17, 104.16, 55.16, 41.95, 39.62, 39.12, 32.67, 31.19, 25.31, 22.79, 22.53. FT-IR (KBr, cm^−1^) 3350.28, 3256.30, 2953.92, 2165.27, 1638.14, 1511.46, 1307.15, 1160.05, 1086.49, 906.70, 820.89, 694.21, 669.69, 543.02. HRMS (+) Calc. for [C_24_H_34_N_5_O_2_]^+^ 423.27125, found: 424.27572 [M+H]^+^. HPLC (protocol A): t_R_ (min) = 9.42. Purity 99.1%.

*3-(tert-Butyl)-N-((S)-1-(((S)-2-(3-chlorophenyl)-1-cyanoethyl)amino)-1-oxo-3-phenylpropan-2-yl)-1-methyl-1H-pyrazole-5-carboxamide (**54**)*

Yield 50%. Yellowish solid. *R_f_* = 0.6 (ethyl acetate: *n*-hexane; 6:4). Mp 132 – 133 °C. ^1^H NMR (200 MHz, CD_3_OD/ CDCl_3_) δ 7.54 – 7.35 (m, 9H), 6.75 (s, 1H), 5.19 – 5.11 (m, 1H), 4.51 – 4.48 (m, 1H), 4.17 (s, 3H), 3.33 – 3.21 (m, 4H), 1.49 (s, 9H). ^13^C NMR (50 MHz, CD_3_OD) δ 172.23, 161.25, 161.03, 137.40, 136.96, 135.83, 135.15, 130.68, 130.07, 129.87, 129.23, 128.42, 128.30, 127.69, 118.18, 104.55, 55.12, 42.41, 38.76, 38.62, 32.51, 30.78. FT-IR (cm^−1^) 3288.99, 2925.31, 2855.85, 2161.18, 1666.74, 1634.05, 1544.15, 1507.38, 1450.17, 1266.29, 1221.34, 1074.23, 861.75, 747.33, 706.47, 694.21. HRMS (+) Calc. for [C_27_H_31_ClN_5_O_2_]^+^ 491.21663, found: 492.21034 [M+H]^+^. HPLC (protocol A): t_R_ (min) = 9.61. Purity > 99.9 %.

*3-(tert-Butyl)-N-((S)-1-(((S)-1-cyano-2-(pyridin-4-yl)ethyl)amino)-1-oxo-3-phenylpropan-2-yl)-1-methyl-1H-pyrazole-5-carboxamide (**55**)*

Yield 35%. Yellowish wax. *R* = 0.4 (ethyl acetate: *n*-hexane; 6:4). ^1^H NMR (400 MHz, CD_3_OD) δ 8.44 (s br, 2H), 7.38 (d, *J* = 5.3 Hz, 2H), 7.30 – 7.22 (m, 5H), 6.63 (s, 1H), 5.14 (t, *J* = 7.5 Hz, 1H), 4.66 (dd, *J* = 8.6, 6.8 Hz, 1H), 3.94 (s, 3H), 3.26 – 3.13 (m, 4H), 1.30 (s, 9H). ^13^C NMR (100 MHz, DMSO-*d_6_*) δ 171.17, 170.90, 159.29, 158.46, 137.89, 136.81, 135.23, 134.81, 129.27, 128.86, 128.13, 127.92, 126.93, 126.34, 118.94, 60.46, 53.87, 53.24, 41.15, 31.39, 30.18, 13.74. FT-IR (cm^−1^) 3264.47, 2953.92, 2913.05, 2851.76, 2161.43, 1662.66, 1605.45, 1548.24, 1466.62, 1204.99, 1115.10, 996.59, 800.45, 735.07, 681.95. HRMS (+) Calc. for [C_26_H_31_N_6_O_2_]^+^ 459.25085, found: 459.25627 [M+H]^+^. HPLC (protocol A): t_R_ (min) = 6.84. Purity 98.6%.

*N-((S)-1-(((1R,2R)-2-(Benzyloxy)-1-cyanopropyl)amino)-1-oxo-3-phenylpropan-2-yl)-3-(tert-butyl)-1-methyl-1H-pyrazole-5-carboxamide (**56**)*

Yield 80%. White solid. *R_f_* = 0.5 (ethyl acetate: *n*-hexane; 6:4). Mp. 115 – 116 °C. ^1^H-NMR (200 MHz, CDCl_3_) δ 7.35 – 7.26 (m, 10H), 6.95 (d, *J* = 8.7 Hz, 1H), 6.61 (d, *J* = 7.0 Hz, 1H), 6.30 (s, 1H), 4.88 – 4.85 (m, 2H), 4.62 – 4.58 (m, 1H), 4.03 (s, 3H), 3.82 – 3. 78 (m, 1H), 3.29 – 3.24 (m, 2H), 1.31 (s, 9H), 1.03 (d, *J* = 6.0 Hz, 3H). ^13^C NMR (50 MHz, CDCl_3_) δ 170.30, 159.83, 159.62, 136.42, 135.27, 133.75, 128.67, 128.41, 128.00, 127.60, 127.30, 126.87, 116.63, 102.65, 72.60, 70.91, 53.85, 44.51, 38.33, 37.60, 31.40, 29.94, 15.29. FT-IR (cm^−1^) 3354.37, 3252.21, 2181.61, 1654.61, 1654.48, 1540.07, 1486.95, 1290.81, 1151.87, 1086.48, 894.44, 808.63, 710.56, 661.32, 543.02. HRMS (+) Calc. for [C_29_H_36_N_5_O_3_]^+^ 502.28182, found: 502.28095 [M+H]^+^. HPLC (protocol A): t_R_ (min) = 9.54. Purity 99.4 %.

*N-((S)-1-(((1S,2S)-2-(Benzyloxy)-1-cyanopropyl)amino)-1-oxo-3-phenylpropan-2-yl)-3-(tert-butyl)-1-methyl-1H-pyrazole-5-carboxamide (**57**)*.

Yield 82%. White solid. *R_f_* = 0.5 (ethyl acetate: *n*-hexane; 6:4). Mp. 123 – 124 °C. ^1^H NMR (200 MHz, CDCl_3_) δ 7.36 – 7.25 (m, 10H), 6.98 (d, *J* = 8.7 Hz, 1H), 6.65 (d, *J* = 7.0 Hz, 1H), 6.32 (s, 1H), 4.91 – 4.83 (m, 2H), 4.63 – 4.60 (m, 1H), 4.05 (s, 3H), 3.80 (s br, 1H), 3.24 – 3.20 (m, 2H), 1.33 (s, 9H), 1.05 (d, *J* = 6.0 Hz, 3H). ^13^C NMR (50 MHz, CDCl_3_) δ 171.01, 160.54, 160.34, 137.14, 135.98, 134.46, 129.39, 129.12, 128.71, 128.32, 128.02, 127.59, 117.34, 103.37, 72.85, 71.15, 54.56, 45.22, 39.04, 38.32, 32.12, 30.65, 16.00. FT-IR (cm^−1^) 2376.73, 2958.00, 2169.35, 1642.22, 15400.07, 1499.21, 1442.00, 1290.81, 1225.43, 1111.01, 1033.37, 988.42, 743.25, 690.12. HRMS (+) Calc. for [C_29_H_36_N_5_O_3_]^+^ 502.28182, found: 502.28095 [M+H]^+^. HPLC (protocol A): t_R_ (min) = 9.43. Purity 98.3 %.

*N-((S)-1-(((1R,2R)-2-(Benzyloxy)-1-cyanopropyl)amino)-4-methyl-1-oxopentan-2-yl)-3-(tert-butyl)-1-methyl-1H-pyrazole-5-carboxamide (**58**)*

Yield 78%. White solid. *R_f_* = 0.5 (ethyl acetate: *n*-hexane; 6:4). Mp. 96 – 97 °C. ^1^H NMR (400 MHz, DMSO-*d_6_*) δ 8.93 (d, *J* = 8.1 Hz, 1H), 8.43 (d, *J* = 7.6 Hz, 1H), 7.38 – 7.29 (m, 5H), 6.88 (s, 1H), 5.07 – 5.04 (m, 1H), 4.64 – 4.61 (m, 2H), 4.59 – 4.56 (m, 1H), 3.97 – 3.94 (m, 3H), 3.86 – 3.84 (m, 1H), 1.72 – 1.64 (m, 2H), 1.50 – 1.46 (m, 2H), 1.26 – 1.20 (m, 12H), 0.93 (d, *J* = 6.3 Hz, 3H), 0.88 (d, *J* = 6.5 Hz, 3H). ^13^C NMR (100 MHz, DMSO-*d_6_*) δ 172.36, 159.56, 158.66, 137.92, 134.92, 128.10, 127.53, 127.45, 117.82, 103.73, 73.39, 70.40, 51.05, 44.75, 38.44, 31.53, 30.31, 24.25, 22.91, 21.19, 15.69. FT-IR (cm^−1^) 3378.89, 3350.28, 3186.83, 2958.00, 2116.43, 1674.91, 1650.40, 1531.90, 1368.44, 1262.20, 1160.05, 1057.89, 1033.37, 780.02, 735.07, 649.26, 604.31. HRMS (+) Calc. for [C_26_H_38_N_5_O_3_] 468.29746, found: 468.29785 [M+H]^+^. HPLC (protocol A): t_R_ (min) = 9.61. Purity 98.3 %.

*N-((S)-1-(((1S,2S)-2-(Benzyloxy)-1-cyanopropyl)amino)-4-methyl-1-oxopentan-2-yl)-3-(tert-butyl)-1-methyl-1H-pyrazole-5-carboxamide (**59**)*

Yield 70%. White solid. *R_f_* = 0.5 (ethyl acetate: *n*-hexane; 6:4). Mp. 110 – 111 °C. ^1^H NMR (400 MHz, CDCl_3_) δ 7.27 – 7.20 (m, 5H), 7.14 (d, *J* = 8.5 Hz, 1H), 6.42 (d, *J* = 7.8 Hz, 1H), 6.33 (s, 1H), 4.85 – 4.82 (m, 1H), 4.62 – 4.52 (m, 3H), 3.96 (s, 3H), 3.87 – 3.84 (m, 1H), 1.69 – 1.59 (m, 3H), 1.22 (s, 9H). 1.18 (d, *J* = 6.3 Hz, 3H) 1.24 – 1.19 (m, 12H), 0.92 – 0.88 (m, 6H). ^13^C NMR (100 MHz, CDCl_3_) δ 172.10, 160.65, 160.51, 137.21, 134.43, 128.66, 128.23, 127.92, 117.45, 103.16, 73.69, 71.80, 51.75, 45.28, 40.81, 38.06, 32.10, 30.62, 25.04, 22.97, 22.13, 16.36. FT-IR (cm^−1^) 3333.94, 3284.90, 2962.09, 2868.10, 2255.16, 1647.44, 1540.07, 1507.38, 1442.00, 1274.46, 1245.86, 988.42, 739.16, 661.52. HRMS (+) Calc. for [C_26_H_38_N_5_O_3_] 468.29746, found: 468.29785 [M+H]^+^. HPLC (protocol A): t_R_ (min) = 9.56. Purity 96.9%. *N-((S)-1-(((1R,2R)-2-(Benzyloxy)-1-cyanopropyl)amino)-3-(3-chlorophenyl)-1-oxopropan-2-yl)-3-(tert-butyl)-1-methyl-1H-pyrazole-5-carboxamide (**60**)*

Yield 50%. Yellowish wax. *R_f_* = 0.4 (ethyl acetate: *n*-hexane; 6:4). ^1^H NMR (400 MHz, CDCl_3_) δ 7.30 – 7.13 (m, 6H), 7.04 (dt, *J* = 7.0, 1.7 Hz, 1H), 6.62 (dd, *J* = 17.8, 8.2 Hz, 2H), 6.25 (s, 1H), 4.77 – 4.71 (m, 2H), 4.48 (dd, *J* = 36.1, 11.7 Hz, 2H), 3.95 (s, 3H), 3.82 – 3.74 (m, 1H), 3.11 – 3.00 (m, 2H), 1.20 (s, 9H), 1.08 (d, *J* = 6.3 Hz, 3H). ^13^C NMR (100 MHz, CDCl_3_) δ 170.73, 160.54, 160.12, 137.80, 136.91, 134.84, 134.30, 130.44, 129.51, 128.71, 128.34, 128.03, 127.84, 127.62, 117.06, 103.35, 73.35, 71.72, 54.28, 45.07, 39.06, 38.13, 32.07, 30.58, 16.21. FT-IR (cm^−1^) 3357.24, 3264.21, 2193.15, 1654.33, 1538.29, 1487.88, 1209.18, 1141.78, 1036.43, 863.33, 807.36, 701.44, 658.23, 534.57. ESI-MS (+) Calc. for [C_29_H_345_ClN_5_O_3_] 536.24284, found: 536.24235 [M+H]^+^.HPLC (protocol B): t_R_ (min) = 12.06. Purity 96.5%.

#### Synthesis of compounds 61-64

Compounds **61**-**64** have been synthesized from compounds **34**-**37** by removal of the benzyl group through hydrogenation on Pd/C (method A).

*N-((S)-1-(((2S,3R)-1-Amino-3-hydroxy-1-oxobutan-2-yl)amino)-1-oxo-3-phenylpropan-2-yl)-3-(tert-butyl)-1-methyl-1H-pyrazole-5-carboxamide.(**61**)*

Yield 92%. Colorless wax. *R_f_* = 0.2 (ethyl acetate). ^1^H NMR (200 MHz, CDCl_3_) δ 8.03 (s, 1H), 7.36 – 6.94 (m, 5H), 6.54 (s, 1H), 4.44 – 4.39 (m, 2H), 4.03 – 4.01 (m, 1H), 3.96 (s, 3H), 3.18 – 3.14 (m, 1H), 2.87 (s, 1H), 1.30 (s, 9H), 1.03 – 1.01 (m, 3H). ^13^C NMR (50 MHz, CDCl_3_) δ 173.54, 172.15, 161.25, 160.35, 136.25, 135.98, 134.58, 129.36, 128.76, 127.25, 66.60, 58.51, 55.55, 54.30, 38.70, 31.98, 30.51, 18.92. ESI-MS (+) Calc. for [C_22_H_31_N_5_O_4_] 429.51, found:452.4 [M+Na]^+^.

*N-((S)-1-(((2R,3S)-1-Amino-3-hydroxy-1-oxobutan-2-yl)amino)-1-oxo-3-phenylpropan-2-yl)-3-(tert-butyl)-1-methyl-1H-pyrazole-5-carboxamide.(**62**)*

Yield 95%. Colorless wax. *R_f_* = 0.2 (ethyl acetate). ^1^H NMR (200 MHz, CDCl_3_) δ 8.01 (s, 1H), 7.36 – 6.98 (m, 5H), 6.59 (s, 1H), 4.42 – 4.36 (m, 2H), 4.04 – 4.01 (m, 1H), 3.96 (s, 3H), 3.12 – 3.09 (m, 1H), 2.89 (s, 1H), 1.33 (s, 9H), 1.03 – 0.99 (m, 3H). ^13^C NMR (50 MHz, CDCl_3_) δ 173.54, 172.15, 161.25, 160.35, 136.25, 135.98, 134.58, 129.36, 128.76, 127.25, 66.60, 58.51, 55.55, 54.30, 38.70, 31.98, 30.51, 18.92. ESI-MS (+) Calc. for [C_22_H_31_N_5_O_4_] 429.51, found: 452.4 [M+Na]^+^.

*N-((S)-1-(((2S,3R)-1-Amino-3-hydroxy-1-oxobutan-2-yl)amino)-4-methyl-1-oxopentan-2-yl)-3-(tert-butyl)-1-methyl-1H-pyrazole-5-carboxamide(**63**)*

Yield 90%. Colorless wax. *R_f_* = 0.2 (ethyl acetate). ^1^H NMR (200 MHz, CDCl_3_) δ 7.37 (d, *J* = 3.9 Hz, 1H), 6.74 – 6.71 (m, 2H), 6.42 (s, 1H), 4.71 – 4.68 (m, 1H), 4.42 – 4.38 (m, 2H), 4.06 (s, 3H), 1.71 – 1.68 (m, 3H), 1.27 (s, 9H), 1.14 – 1.11 (m, 3H), 0.98 – 0.92 (m, 6H). ESI-MS (+) Calc. for [C_19_H_33_N_5_O_4_] 395.51, found: 418.5 [M+Na]^+^.

*N-((S)-1-(((2R,3S)-1-Amino-3-hydroxy-1-oxobutan-2-yl)amino)-4-methyl-1-oxopentan-2-yl)-3-(tert-butyl)-1-methyl-1H-pyrazole-5-carboxamide (**64**)*

Yield 93%. Colorless wax. *R_f_* = 0.2 (ethyl acetate). ^1^H NMR (200 MHz, CDCl_3_) δ 7.34 (d, *J* = 3.9 Hz, 1H), 6.79 – 6.75 (m, 2H), 6.46 (s, 1H), 4.73 – 4.70(m, 1H), 4.45 – 4.41 (m, 2H), 4.11 (s, 3H), 1.77 – 1.74 (m, 3H), 1.28 (s, 9H), 1.19 – 1.17 (m, 3H), 0.99 – 0.91 (m, 6H). ESI-MS (+) Calc. for [C_19_H_33_N_5_O_4_] 395.51, found: 418.5 [M+Na]^+^.

#### Synthesis of compounds 65-68

Compounds **65**-**68** have been synthesized by dehydration of the corresponding primary amide precursor **61**-**64** with trifluoroacetic anhydride (method B).

*3-(tert-Butyl)-N-((S)-1-(((1R,2R)-1-cyano-2-hydroxypropyl)amino)-1-oxo-3-phenylpropan-2-yl)-1-methyl-1H-pyrazole-5-carboxamide (**65**)*

Yield 45%. White solid. *R_f_* = 0.7(ethyl acetate). Mp. 159 – 160 °C. ^1^H NMR (500 MHz, CDCl_3_) δ 7.32 – 7.26 (m, 4H), 7.26 – 7.21 (m, 1H), 6.65 (s, 1H), 4.84 – 4.79 (m, 2H), 3.96 (s, 3H), 3.88 – 3.84 (m, 1H), 3.20 (dd, *J* = 13.6, 7.1 Hz, 1H), 3.10 (dd, *J* = 13.6, 7.1 Hz, 1H), 1.31 (s, 9H), 1.07 (d, *J* = 6.3 Hz, 3H). ^13^C NMR (125 MHz, CDCl_3_) δ 173.35, 162.06, 161.60, 138.15, 136.69, 130.42, 129.57, 127.98, 118.24, 105.04, 67.67, 56.05, 38.86, 38.68, 32.90, 30.87, 18.89, 18.55. FT-IR (KBr, cm^−1^) 3293.08, 2953.92, 2913.05, 2868.10, 1646.10, 1540.07, 1446.08, 1286.72, 1237.68, 1098.75, 922.51, 739.16, 689.30, 469.47. HRMS (+) Calc. for [C_22_H_30_N_5_O_3_]^+^ 412.23486, found: 412.23811 [M+H]^+^. HPLC (protocol A): t_R_ (min) = 7.65. Purity 97.8%.

*3-(tert-Butyl)-N-((S)-1-(((1S,2S)-1-cyano-2-hydroxypropyl)amino)-1-oxo-3-phenylpropan-2-yl)-1-methyl-1H-pyrazole-5-carboxamide (**66**)*

Yield 42%. White solid. *R_f_* = 0.7 (ethyl acetate). Mp. 145 – 147 °C. ^1^H NMR (400 MHz, CDCl_3_) δ 7.31 – 7.23 (m, 5H), 7.07 (d, *J* = 8.0 Hz, 1H), 6.67 (d, *J* = 7.5 Hz, 1H), 6.31 (s, 1H), 4.85 – 4.84 (m, 1H), 4.72 – 4.68 (m, 1H), 4.15 – 4.10 (m, 1H), 4.02 (s, 3H), 3.21 – 3.15 (m, 2H), 1.29 – 1.27 (m, 9H), 1.08 (d, *J* = 6.2 Hz, 3H). ^13^C NMR (100 MHz, CDCl_3_) δ 172.41, 171.26, 160.58, 135.89, 134.27, 129.38, 129.14, 127.65, 117.25, 103.52, 67.39, 54.76, 47.02, 39.11, 38.38, 32.09, 30.59, 19.02. FT-IR (KBr, cm^−1^) 3289.11, 2955.43, 2918.55, 2860.11, 1649.54, 1543.16, 1444.02, 1277.62, 1233.66, 1091.87, 944.77, 745.55, 690.34, 477.11. HRMS (+) Calc. for [C_22_H_30_N_5_O_3_]^+^ 412.23486, found: 412.23811 [M+H]^+^.HPLC (protocol A): t_R_ (min) = 7.71. Purity 95.7%.

*3-(tert-Butyl)-N-((S)-1-(((1R,2R)-1-cyano-2-hydroxypropyl)amino)-4-methyl-1-oxopentan-2-yl)-1-methyl-1H-pyrazole-5-carboxamide (**67**)*

Yield 33%. White solid. *R_f_* = 0.5 (ethyl acetate). Mp. 118 – 119 °C. ^1^H NMR (400 MHz, CDCl_3_) δ 7.56 – 7.51 (m, 1H), 6.80 – 6.75 (m, 1H), 6.43 (s, 1H), 4.87 – 4.83 (m, 1H), 4.65 – 4.63 (m, 1H), 4.15 – 4.11 (m, 1H), 4.06 (s, 3H), 1.31 – 1.20 (m, 15H), 0.97 (2d, *J* = 6.2 Hz, 3H). ^13^C NMR (100 MHz, CDCl_3_) δ 174.32, 161.62, 162.14, 135.77, 117.22, 104.56, 67.34, 52.68, 41.12, 38.28, 31.46, 32.15, 26.11, 23.56, 21.65, 19.03. FT-IR (KBr, cm^−1^) 3297.16, 2953.92, 2913.05, 2868.10, 1646.31, 1535.89, 1503.29, 1462.52, 1364.36, 1282.63, 1172.38, 1131.44, 996.59, 730.99, 592.05. HRMS (+) Calc. for [C_19_H_32_N_5_O_3_]^+^ 378.25071, found: 378.25231 [M+H]^+^.HPLC (protocol A): t_R_ (min) = 8.08. Purity: 96.7%.

*3-(tert-Butyl)-N-((S)-1-(((1S,2S)-1-cyano-2-hydroxypropyl)amino)-4-methyl-1-oxopentan-2-yl)-1-methyl-1H-pyrazole-5-carboxamide (**68**)*

Yield 35%. White solid. *R_f_* = 0.5 (ethyl acetate). Mp. 111 – 113 °C. ^1^H NMR (400 MHz, CD_3_OD) δ 6.78 (s, 1H), 4.88 – 4.84 (m, 1H), 4.65 – 4.60 (m, 1H), 4.07 – 4.05 (m, 1H), 4.05 (s, 3H), 1.81 – 1.61 (m, 4H), 1.32 (s, 9H), 1.26 (d, *J* = 6.3 Hz, 3H), 1.00 (2d, *J* = 6.3 Hz, 6H). ^13^C NMR (100 MHz,CD_3_OD) δ 175.01, 162.46, 161.76, 136.76, 118.59, 105.26, 68.00, 53.28, 41.76, 39.00, 33.07, 31.04, 26.27, 23.54, 21.94, 19.44. FT-IR (KBr, cm^−1^) 3293.08, 2962.09, 2917.14, 2847.67, 1646.31, 1540.07, 1458.34, 1278.55, 1131.44, 1078.32, 1000.68, 853.58, 812.71, 780.02, 730.99, 559.36. HRMS (+) Calc. for [C_19_H_32_N_5_O_3_]^+^ 378.25071, found: 378.25231 [M+H]^+^. HPLC (protocol A): t_R_ (min) = 7.94. Purity: 98.6%.

#### Synthesis of compound 69

Compound **69** has been synthesized by removal of the benzyl group from compound **60** with DDQ (method B).

*3-(tert-butyl)-N-((S)-3-(3-chlorophenyl)-1-(((1R,2R)-1-cyano-2-hydroxypropyl)amino)-1-oxopropan-2-yl)-1-methyl-1H-pyrazole-5-carboxamide (**69**)*

Yield 32%. White solid. *R_f_* = 0.3 (ethyl acetate). Mp. 102 – 103 °C. ^1^H NMR (400 MHz, CD_3_OD) δ 7.35 – 7.28 (m, 4H), 6.69 (s, 1H), 4.87 – 4.83 (m, 2H), 3.99 (s, 3H), 3.90 – 3.84 (m, 1H), 3.24 (dd, *J* = 13.6, 7.1 Hz, 1H), 3.08 (dd, *J* = 13.6, 6.5 Hz, 1H), 1.34 (s, 9H), 1.11 (d, *J* = 6.3 Hz, 3H). ^13^C NMR (100 MHz, CD_3_OD) δ 175.23, 163.23, 151.94, 151.48, 128.04, 126.57, 120.30, 120.10, 119.52, 119.45, 117.86, 108.12, 94.92, 57.55, 45.93, 28.74, 28.56, 22.78, 20.75, 18.78. FT-IR (KBr, cm^−1^) 3284.90, 3056.07, 2958.00, 2925.31, 2868.10, 1638.14, 1556.41, 1499.21, 1462.43, 1364.36, 1270.37, 1229.51, 1098.75, 878.09, 730.99, 702.38, 453.12. ESI-MS (+) Calc. for [C_22_H_29_ClN_5_O_3_]^+^ 446.19589, found: 446.19745 [M+H]^+^.HPLC (protocol A): t_R_ (min) = 8.55. Purity 99.7%.

### Enzyme inhibition studies

Enzyme inhibition studies for Cz and LmCPB were performed as previously described for Cz [14]. Cathepsins B, L, S were assayed as reported [17, 18].

Cathepsin K assay. Human recombinant cathepsin K was assayed on a FLUOSTAR Optima plate reader at 25 °C with an excitation wavelength of 360 nm and an emission wavelength of 440 nm on a 96 well plate. The enzyme solution (23 µg/mL in 50 mM sodium acetate pH 5.5, 50 mM NaCl, 0.5 mM EDTA, 5 mM DTT) was diluted 1:100 with assay buffer (100 mM sodium citrate buffer pH 5.0, 100 mM NaCl, 1 mM EDTA, 0.01% CHAPS) containing 5 mM DTT and was then incubated at 37 °C for 30 min for activation. A 1.5 mM stock solution of the substrate Z-Leu-Arg-AMC was prepared in DMSO. The final substrate concentration was 6 µM (= 3.05 × *K*_m_). The assay was performed with a final concentration of cathepsin K of 1.73 ng/mL. Stock solutions of inhibitors were prepared in DMSO. The final DMSO concentration was 2% (4 µL). Into a well containing 194.5 µL assay buffer, 0.8 µL of the fluorogenic substrate, DMSO and inhibitor solution (3.2 µL) were added. Upon addition of cathepsin K (1.5 µL), the measurement was started and followed for 20 min.

### In vitro trypanocidal activity evaluation on intracellular amastigote forms (Tulahuen strain)

Cells were analyzed in 96-well plates, cells from the LLCMK_2_ strain were plated at a concentration of 5×10^4^ cells/mL. Trypomastigote forms of the Tulahuen LacZ strain were added at a concentration of 5×10^5^ parasites mL^−1^ and placed in the incubator at 37 °C with 5% CO_2_ for 48 hours. After the incubation period, the trypomastigote forms present were removed by successive washes with PBS, remaining only as intracellular amastigote forms. Compounds were added at different concentrations (1.95 μM to 250 μM serial dilutions) and incubated for 72 hours. At the end of this period, the substrate CPRG (chlorophenol red β-D-galactopyranoside, 400 μM in 0.3% Triton X-100, pH 7.4) was added. After 4 hours of incubation at 37 °C, the plates were analyzed in a spectrophotometer at 570 nm to obtain the effective concentration (EC_50_) to reduce the parasitemia inside the host cell. Benznidazole was used as a positive control in the same concentrations as the substances, and DMSO as a negative control. Compounds were solubilized in DMSO. The same assay condition was performed to determine the cytotoxic concentration data (CC_50_) using the non-infected host cells. The selectivity index (SI) was calculated using the formula: SI = EC_50_/CC_50_. All statistical analyses were done with the program GraphPad Prism v.5.

## Results

### Structure-based design, modelling studies, and compound synthesis

Cz, the recombinant form of cruzipain is a monomeric enzyme, composed of two folded and equally sized domains. These domains are divided by the enzyme’s active site, which is V-shaped and largely exposed to solvent. A catalytic triad cysteine-histidine-asparagine forms the active site [13]. The main polar interactions between the protein and inhibitor are well conserved involving the residues Gln19, Gly66, Asp161, His162, and Trp184 of the enzyme. Cz is a cathepsin L-like cysteine protease and is closely related to the mammalian CPs such as CatB, CatK, CatL, and CatS.

A variety of studies have been conducted on optimization strategies for the interactions of different classes of inhibitors with the S1, S2 and S3 binding sites of cruzain and related cysteine proteases [13,14,19,20]. Nonetheless, far less is known about the attainable interactions at S1′ for dipeptidyl nitrile inhibitors [21]. The high-resolution crystal structure of cruzain shows that there is a large open surface characterized by Trp177 in the primed binding site region (Fig 3) [22]. The design of compounds to exploit this cavity would provide enhanced enzyme-inhibitor interactions. This concept has been already applied for a class of different dipeptidic vinyl sulfone inhibitors [23]. As Fig 3 (left) exemplarily illustrates, the substituents of vinyl sulfone inhibitors predominantly sit on top of the shelf formed by residues Ser139, Met142 and Asp158 rather than adopting an orientation for a strong aromatic–aromatic interactions with Trp177. The nitrile inhibitor 33L does not bear an appropriate substituent that would allow for an interaction with the primed binding region of cruzain (Fig 3, right).

**Fig 3.**
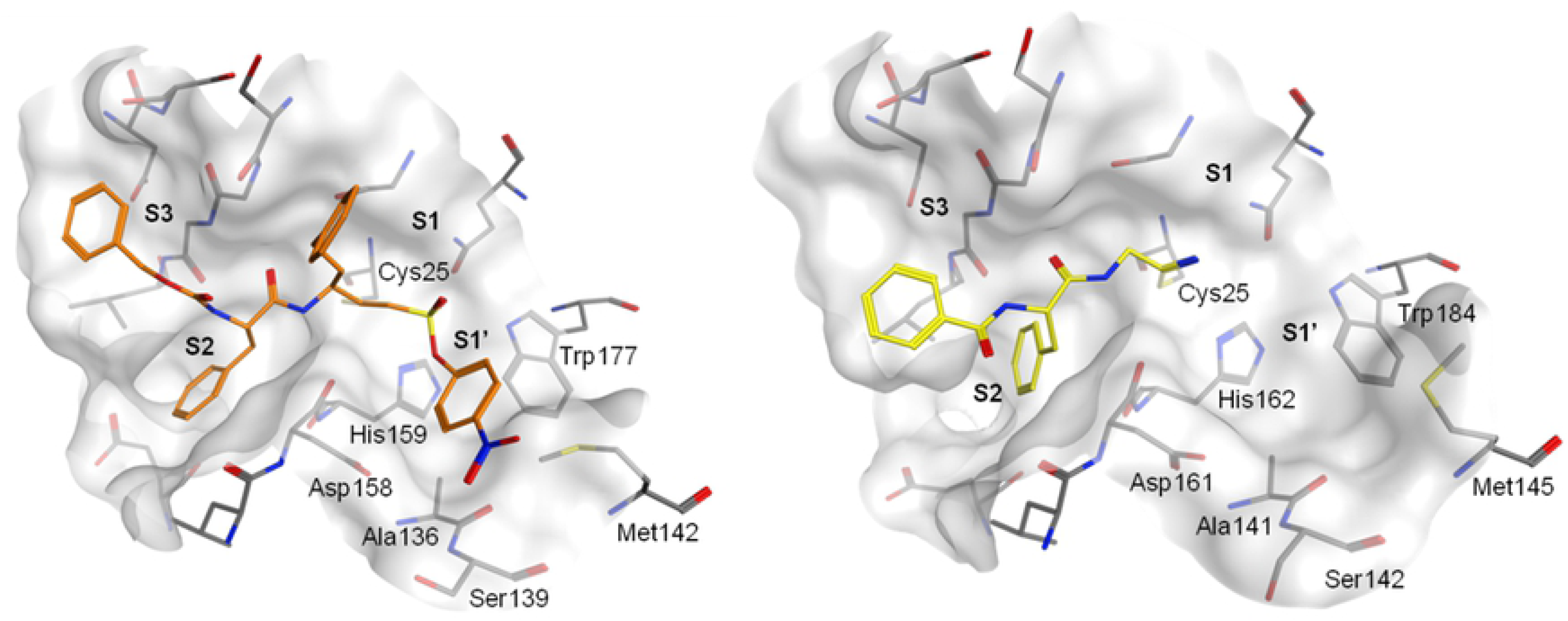
Crystal structures of vinyl sulfone derivative K777 and dipeptidyl nitrile 33L covalently bound to cruzain. Left picture, PDB-ID: 1F2B; right picture, PDB-ID 4QH6.

The nitrile warhead has been applied successfully for a variety of series of cathepsin inhibitors. Peptidic nitriles are known the interact with the active site cysteine by forming a covalent, but reversible thioimidate adduct [24]. The nitrile warhead was also repurposed for Cz inhibition as trypanocidal agents, displaying low toxicity, probably due to the reversible character of interaction [25]. Therefore, starting from our recent study on dipeptidyl nitriles as trypanocidal agents, we expanded our previous inhibitor series to map the S1/S1′ subsites of Cz [13]. By applying a knowledge-based design approach, we have explored different amino acids as possible building blocks for the P1 moiety. Based on a template crystal structure of the dipeptidyl nitrile inhibitor 33L bound to Cz (PDB ID: 4QH6), structural modifications have been executed that might increase the affinity towards the S1’ specificity pocket. Fig 4 shows dipeptidyl nitriles **50**, **52**, **56**, and **58** with different lipophilic substitution patterns at the P1 position, which were assumed to accommodate the S1′ pocket through hydrophobic interactions without interfering in the general mode of binding.

**Fig 4.**
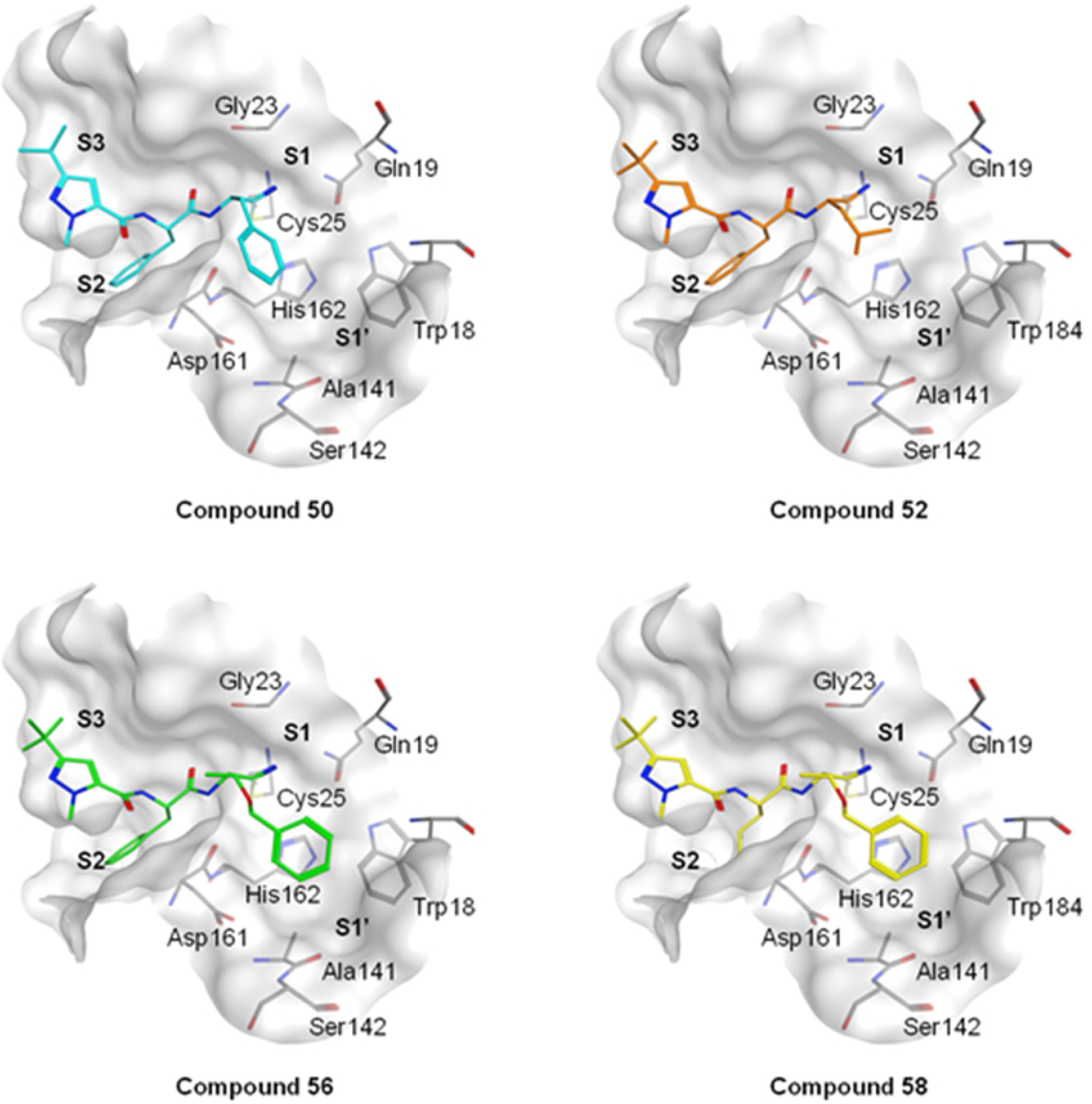
The putative orientation of P1 moieties in compounds 50, 52, 56, and 58. Possible interactions with residues forming pockets S1 and S1’ (PDB ID: 4QH6).

Compound **9** (Fig 5, Fig 6) was adopted as an archetype, with the cyclopropyl group at P1 position and phenylalanine as well as a pyrazole moiety for advantageous interactions with the non-primed binding region of the target protease. We mainly used different natural and unnatural amino acids for the P1 moiety and maintained the nitrile warhead. Leucine (Leu) and phenylalanine (Phe) were incorporated (compounds **50** – **54**, Fig 6) as molecular sensors for aliphatic and aromatic interactions. 4-Pyridylalanine was implemented to leverage the affinity by polar interaction with Asp161. Thr-O-Bzl, an unusual building block for peptide inhibitors, was used as chimera for aliphatic and aromatic interactions. After removal of the benzyl protecting group from Thr-O-Bzl, the so produced alcoholic moiety should allow to evaluating whether a hydrogen bond donor is tolerated in the S1/S1′ area. Moreover, it was intended to investigate how the stereochemistry in this region will influence the affinity with Cz.

**Fig 5.**
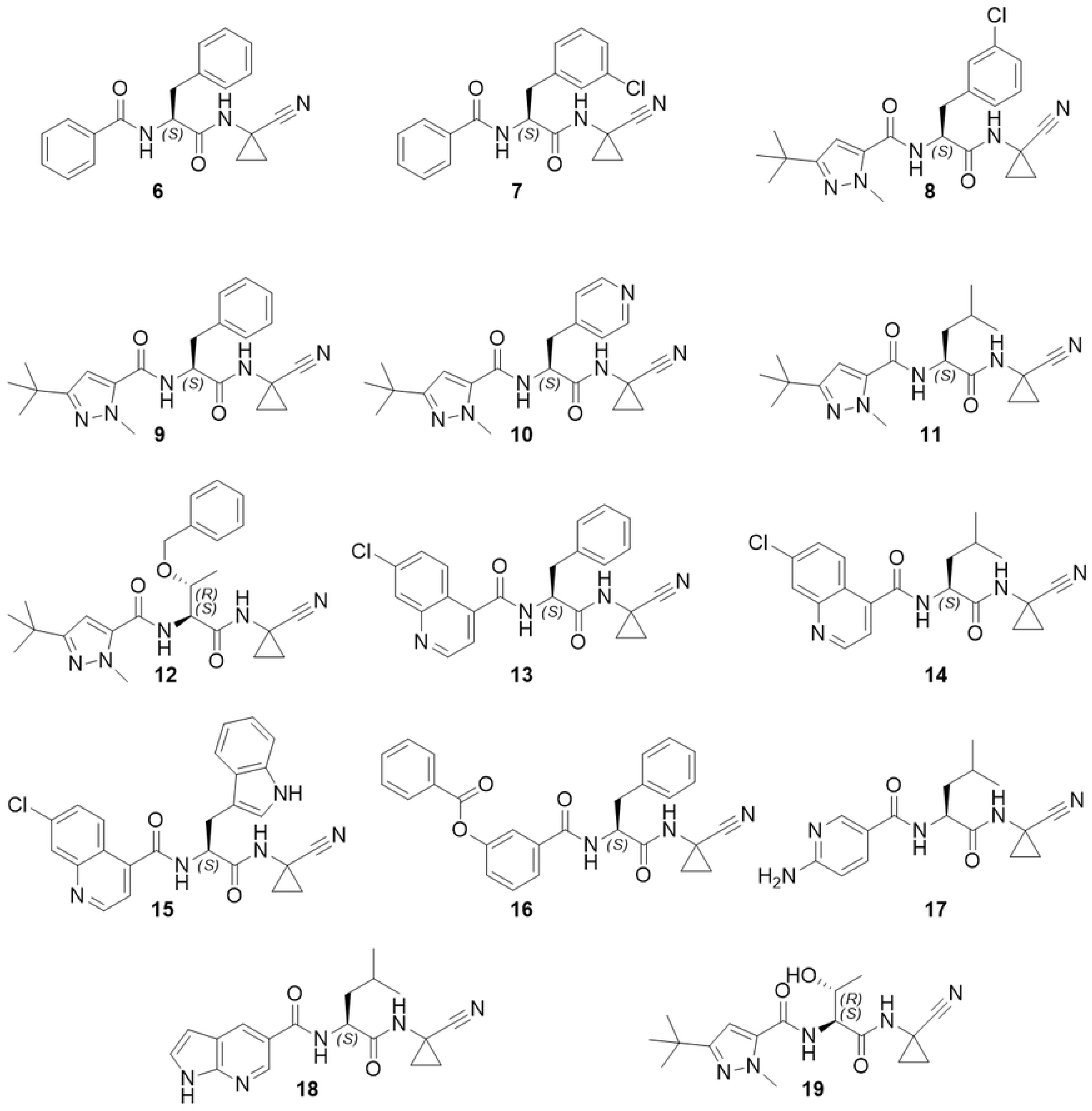
**Structure representation of compounds 6-19.**

**Fig 6.**
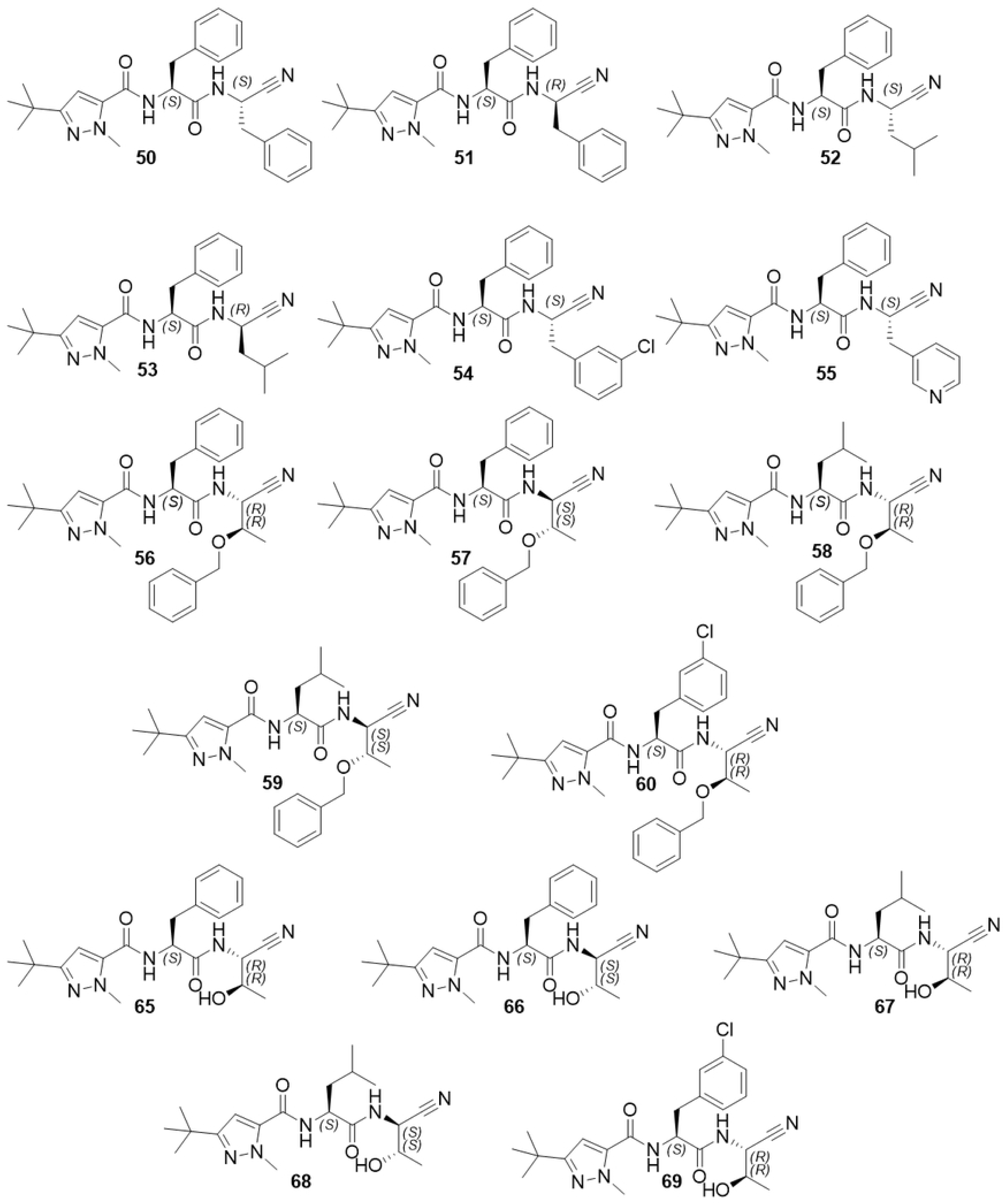
**Structure of compounds 50-60 and 65-69.**

3-(*tert*-Butyl)-1-methyl-1*H*-pyrazole-5-carboxylic acid is a privileged building block applied for the inhibition of Cz and CatL [26]. Thus, we have explored some possible bioisosteres in order to increase the affinity and the selectivity towards Cz, such as 7-chloroquinoline carboxylic acid, 1*H*-indole-5-carboxylic acid, or 6-aminonicotinic acid.

Accordingly, we synthesized a new series of dipeptidyl nitriles (Fig 1, Fig 2). For compounds **6**-**12** and **14**-**19** bearing a cyclopropyl moiety in P1, the synthesis was carried out as known from the literature [14]. The peptide coupling reaction was performed twice; first to connect the enantiomerically pure, Boc-protected P2 amino acid with the aminonitrile moiety, and secondly, after removing the Boc group, to introduce the corresponding aroyl acids (Fig 1). Compound **19** was synthesized from compound **12** by removing the benzyl group under mild oxidative conditions (Fig 1) [27].

For the synthesis of compounds **50**-**60** (Fig. 6), we have adopted a different synthetic strategy. In general, the desired dipeptidyl primary amide was synthesized, followed by the dehydration reaction to form the dipeptidyl nitrile. Due to the diversity of building blocks, it was necessary to evaluate different dehydrating reagents, aiming at the best yield and prevention of racemization. For compounds **65**-**68** the cleavage of the benzyl group was performed by hydrogenolysis before the conversion of the primary amide to the nitrile, while for compound **69**, considering the lability of the chlorine atom under hydrogenolysis, we first transformed the primary amide to the nitrile and then removed the benzyl group under mild oxidative conditions [27]. The absolute geometry of the P1 group did not change, but, owing to CIP priority rules, the configuration at the α−carbon for the Thr-O-Bzl building block changed two times: (i) in the dehydration step to form the nitrile group and, (ii) when the catalytic cysteine attacks the carbon atom of the nitrile warhead to form a covalent bond (Fig 7).

**Fig 7.**
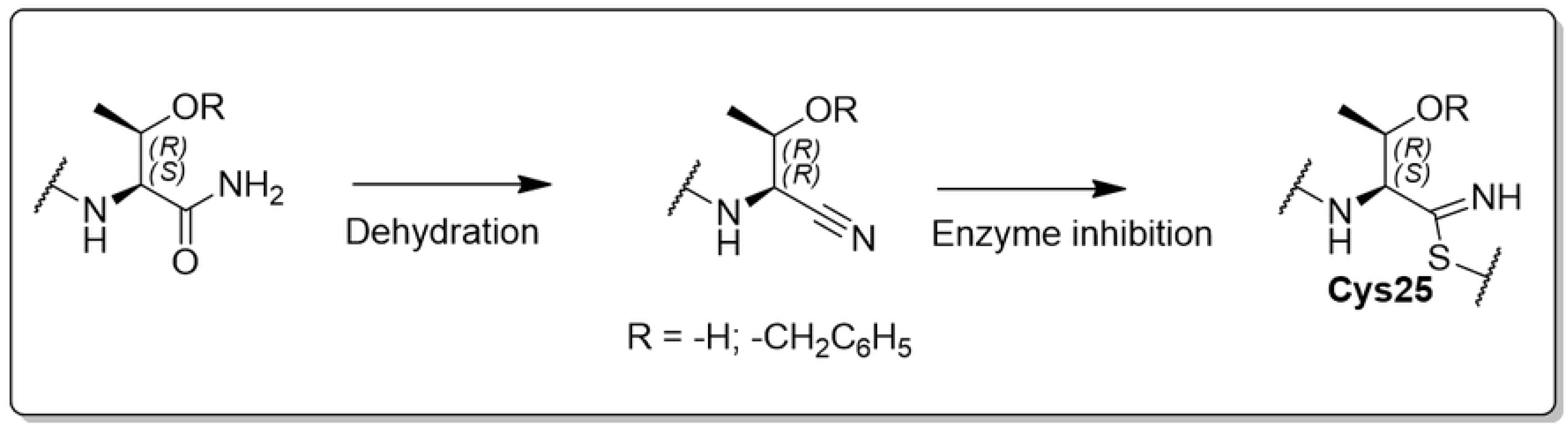
**Change in stereochemistry for compounds bearing Thr or Thr-O-Bzl group in P1.**

### Structure-activity relationships for inhibition of cysteine proteases by dipeptidyl nitriles

The p*K*_i_ values were determined for parasite cysteine proteases (Cz, LmCPB) and also for human cysteine cathepsins (CatB, CatK, CatL, CatS) and are reported in Table 1. Compounds **6**, **8**, **9** and **11** have already been described as competitive inhibitors that bind reversibly to Cz [13]. Some of the compounds are Cz nanomolar inhibitors and also exhibit good affinity for LmCPB, CatL, and CatK. The application of such inhibitors extends to candidates for antiprotozoal action and also as inhibitors of cysteine cathepsins of human host cells in various pathological conditions.

**Table 1.** Structures representation, number identification, p*K*_i_ values for CatB, CatK, CatL, CatS, Cz and LmCPB.

| Cmpd. | $pK_i$ values or remaining activity (%) at 10 μM<sup>a,b</sup> | | | | | |
| --- | --- | --- | --- | --- | --- | --- |
|  | CatB | CatK | CatL | CatS | Cz | LmCPB |
| **6** | 73%<sup>b</sup> | 6.4 | 7.4 | 7.1 | 6.6 | 6.6 |
| **7** | 5.5 | 6.2 | 7.3 | 7.7 | 6.9 | 6.2 |
| **8** | 5.4 | 6.4 | 8.6 | 6.7 | 7.4 | 6.7 |
| **9** | 4.8 | 6.5 | 8.2 | 6.8 | 7.3 | 7.1 |
| **10** | 93%<sup>b</sup> | 5.0 | 7.6 | 5.6 | 6.6 | 6.4 |
| **11** | 4.5 | 8.3 | 7.6 | 7.4 | 7.8 | 7.3 |
| **12** | 4.4 | 96%<sup>b</sup> | 5.9 | 5.9 | 5.1 | 5.5 |
| **13** | 4.9 | 81%<sup>b</sup> | 7.2 | 6.7 | 6.7 | 6.9 |
| **14** | 89%<sup>b</sup> | 6.9 | 6.6 | 6.9 | 7.8 | 7.3 |
| **15** | 4.7 | n.i.<sup>c</sup> | 5.7 | 6.0 | 6.2 | 5.7 |
| **16** | 5.2 | 6.2 | 6.3 | 7.3 | 6.9 | 7.0 |
| **17** | n.i.<sup>c</sup> | 8.0 | 6.5 | 7.6 | 7.1 | 6.8 |
| **18** | 4.5 | 8.7 | 7.1 | 7.3 | 7.7 | 7.2 |
| **19** | 4.6 | 5.5 | 5.2 | 5.0 | 51%<sup>b</sup> | 67%<sup>b</sup> |
| **50** | 4.5 | 6.3 | 8.5 | 7.3 | 7.7 | 7.8 |
| **51** | 94%<sup>b</sup> | 5.6 | 6.9 | 5.9 | 6.3 | 6.1 |
| **52** | 5.1 | 6.7 | 8.3 | 6.9 | 7.5 | 7.4 |
| **53** | 90%<sup>b</sup> | 6.0 | 7.0 | 6.0 | 6.5 | 6.4 |
| **54** | 4.9 | 6.3 | 8.3 | 7.1 | 7.5 | 7.5 |
| **55** | 4.8 | 6.5 | 8.3 | 7.2 | 7.5 | 7.2 |
| **56** | 5.2 | 5.7 | 7.0 | 6.0 | 6.3 | 6.5 |
| **57** | n.i.<sup>c</sup> | 97%<sup>b</sup> | 85%<sup>b</sup> | n.i.<sup>c</sup> | 75%<sup>b</sup> | 72%<sup>b</sup> |
| **58** | 5.1 | 7.8 | 7.2 | 7.3 | 7.9 | 7.7 |
| **59** | n.i.<sup>c</sup> | n.i.<sup>c</sup> | 90%<sup>b</sup> | 85%<sup>b</sup> | 60%<sup>b</sup> | 75%<sup>b</sup> |
| **60** | 6.3 | 6.0 | 8.1 | 6.5 | 7.2 | 6.8 |
| **65** | 92%<sup>b</sup> | 86%<sup>b</sup> | 5.3 | 5.3 | 68%<sup>b</sup> | 69%<sup>b</sup> |
| **66** | 89%<sup>b</sup> | 81%<sup>b</sup> | 5.6 | 4.7 | 5.4 | 4.8 |
| **67** | 92%<sup>b</sup> | 7.8 | 7.0 | 5.0 | 7.1 | 6.7 |
| **68** | 93%<sup>b</sup> | 5.1 | 88%<sup>b</sup> | 83%<sup>b</sup> | 5.4 | 54%<sup>b</sup> |
| **69** | 5.2 | 6.1 | 8.2 | 6.6 | 6.9 | 6.0 |

<sup>a</sup> The standard deviation was lower than 15% for all reported $pK_i$ values. <sup>b</sup> Percentage of remaining activity at 10 μM, n = 2. <sup>c</sup> n.i. = no inhibition observed at 10 μM.

One important question is the cross-reactivity of CP inhibitors, for which an extrathermodynamic relationships can be formulated [28, 29]. The nature of the ligand-target interaction governed by the thermodynamic parameter of the free energy change (via the estimation of the dissociation constant) results in respective extrathermodynamic relationships for a set of derivatives. So, we investigated the degree of linear correlation between Cz and the other CPs by plotting the p*K*_i_ data against each other. The results (see SI) indicated an extrathermodynamic relationship between Cz and LmCPB, while this was not observed for all the other CPs. This finding highlight that the mode of inhibition for this series of compounds is similar for Cz and LmCPB, corroborating the fact that all the structural transformations of prototype compounds **9** and **11** affected the affinity towards the two protozoa CPs with the same magnitude (Fig 8, Fig 9).

**Fig 8.**
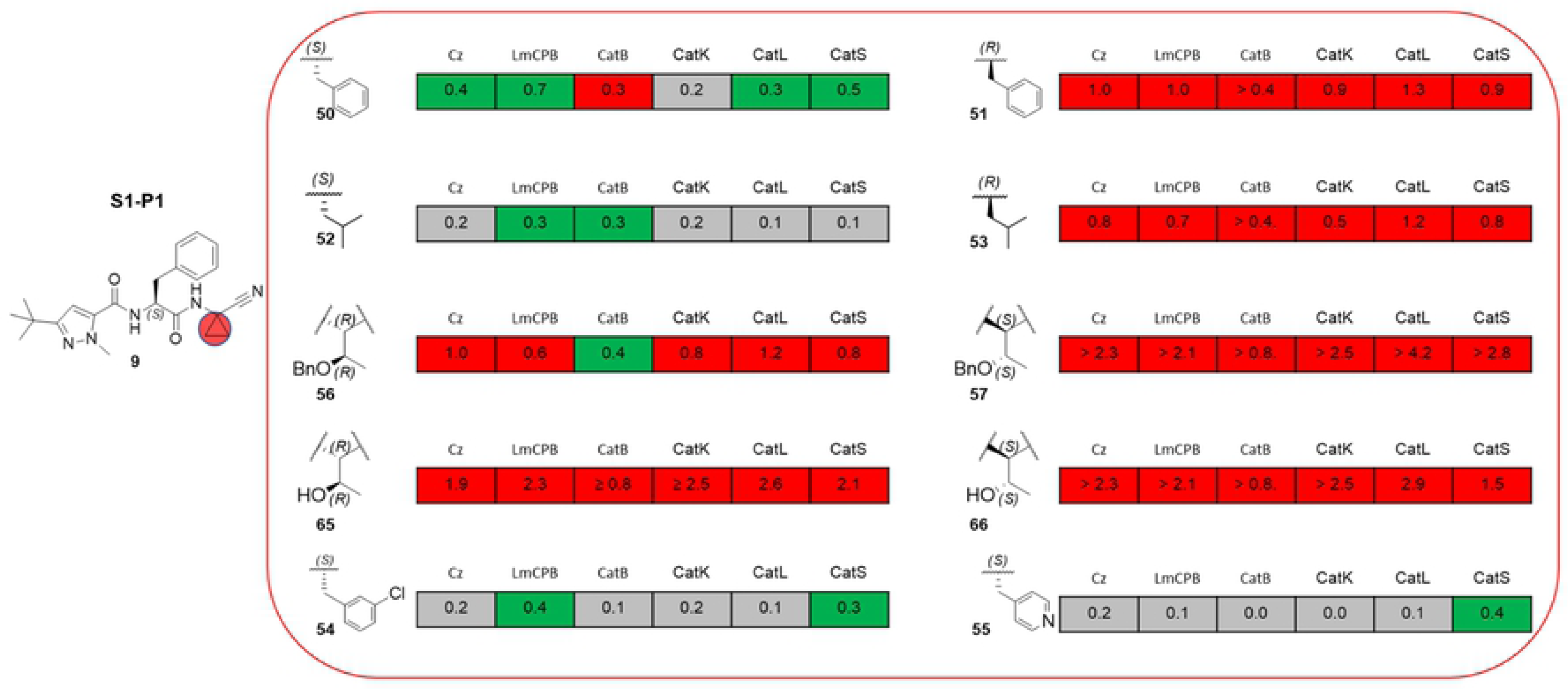
SAR summary for S1-P1 interactions. Values are reported as differences in p*K*_i_ and are color-coded as red (negative), green (positive), grey (no significant difference, Δp*K*_i_ < 0.2).

**Fig 9.**
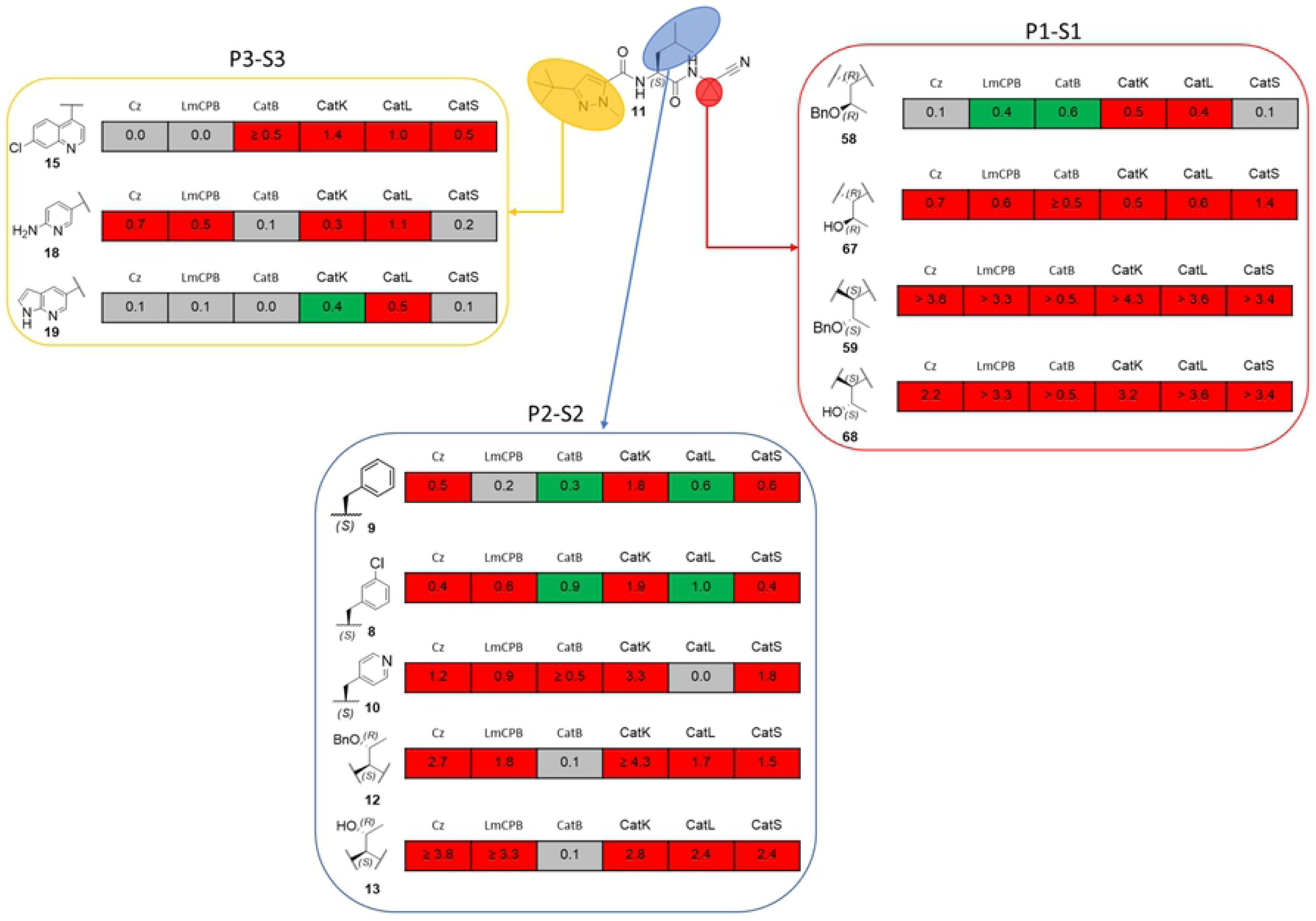
SAR summary starting from compound 11. Values are reported as differences in p*K*_i_ and are color-coded as red (negative), green (positive), grey (no significant difference, Δp*K*_i_ < 0.2).

One common approach to SAR analysis is to examine Δp*K*_i_ values associated with particular structural transformations, and these can be specified concisely using the square bracket notation previously described [30]. For example, the structural transformation of the phenylalanine in P2 of compound **6** to the corresponding 3-chloro-phenylalanine (**8**) can be noted as [**6**→**8**]. As already described for Cz [13], the exchange of benzoyl by 1-methyl-3-*tert*-butyl-pyrazolyl-carbonyl [**6**→**9**] led to a potency increase of 0.5 log units. Following this path, we inserted a meta-benzoic ester in the P3 position in the attempt to design a prodrug analogue [14]. The transformation [**6**→**16**] unfortunately displayed a slight increment as compared to the transformation [**6**→**14**]. Hence, compound **9** (p*K*_i_ of 7.4 for Cz) has been used as a prototype for mapping S1-P1 interaction on different targets (Fig 8).

The effects on affinity resulting from stereochemical modifications in P1 of the structural prototype **9** are shown in Fig 8. In general, the stereochemistry of P1 moiety strongly influences the affinity towards all the CPs. The (*S*)→(*R*) conversion in [**50**→**51**] and [**52**→**53**] decrease the p*K*_i_ values for Cz and LmCP both by about one log unit. Likewise, the double stereochemical modification from (*R*,*R*) benzyl-protected threonine to the (*S*,*S*) diastereomer [**56**→**57**] led to a complete all-target affinity loss. Instead, the structural transformation from the cyclopropyl unit to CH attached with a benzyl group [**9**→**50**] resulted in a significant affinity increment for Cz and LmCPB of 0.4 and 0.7 log units, respectively. Replacement of the P1 cyclopropane linker with CH attached to isopropyl [**9**→**52**], 3-chlorobenzyl [**9**→**54**], or even 4-pyridyl [**9**→**55**] led to a small increase or no significant difference in affinity against those two protozoa enzymes. Moreover, the insertion of the benzyl protected threonine in P1 [**9**→**56**] and [**9**→**57**] decreased the affinity for Cz and LmCPB. Remarkably, replacement by the hydroxybutyl residues led to an almost one hundredfold affinity loss for both enzymes. Essentially, the same trend in affinity was observed for the four mammalian CPs, when the structural modifications in P1 were realized starting from the prototype compound **9** as illustrated in Fig 6. Singularly, the introduction of a benzyl-protected threonine [**9**→**50**] resulted in an *p*K*_i_* decrease of 0.4 log units towards CatB.

As recently described [13], the effects on affinity when replacing the P2 phenylalanine (**9**) with leucine (**11**) appears to depend on the substructural context, and this relates to non-additivity in the SAR. Accordingly, we used compound **11** as a starting prototype for another SAR considering P1, P2, and P3 for structural modifications as summarized in Fig 9.

Substitution of the in P3 positioned 1-methyl-3-*tert*-butyl-pyrazole ring with 7-chloro-quinoline (**15**), or 1*H*-indole (**18**) preserved the high affinity towards Cz and LmCPB, and, strikingly, this substitution led to a decrease of 1.0 and 0.5 in the p*K*_i_ value for CatL (Fig 9). Noteworthy, when the 7-chloro-quinoline moiety was retained in P3 and leucine was exchanged for tryptophan in P2 [**14**→**15**], a huge loss in affinity and selectivity was observed. Insertion of a basic moiety, *i.e.*, 2-amino pyridine [**11**→**17**], produced a significant reduction of potency for Cz (−0.7) and LmCPB (−0.5). Compound **18** showed a high affinity for CatK (p*K*_i_ of 8.7) with a significant selectivity over the other mammalian CPs (Fig 9). In P2 position, the transformation of the leucine moiety to phenylalanine or its derivatives resulted in a loss in affinity up to one log unit for Cz. A similar replacement led to a gain in affinity towards CatL and CatB, and it is consistent with previously reported data [31]. For compound **11**, as for compound **9**, the stereochemistry of the substituent in P1 was vitally for the bimolecular recognition process. Moreover, the transformation [**11**→**58**] kept the p*K*_i_ in the same range for Cz while increasing it by 0.4 log units for LmCPB.

Non-additivity in SAR is of considerable interest [32], and this is illustrated for the six cysteine proteases in Fig 10. Non-additivity can be quantified by comparing the Δp*K*_i_ value resulting from a pair of substructural transformations with the sum of Δp*K*_i_ values that result from the individual transformations. The Cz Δp*K*_i_ values for [**9**→**11**] (0.5) and [**9**→**56**] (−1.0) shown in Fig 10A sum up to −0.5. Nevertheless [**9**→**58**], which corresponds the simultaneous application of the pair transformations, is associated with a Δp*K*_i_ value increase of +0.6, thus indicating that the effects of this pair of transformations on Cz affinity are superadditive. The same was true for the effects of these two transformations on the other five CPs. Analogous analysis of the results in Fig 10B displays the effects of two transformations to be superadditive for the entire CP targets investigated herein. These results entail the P1-S1 and P1-S1’ interactions to be driven by the molecular recognition in P2.

**Fig 10.**
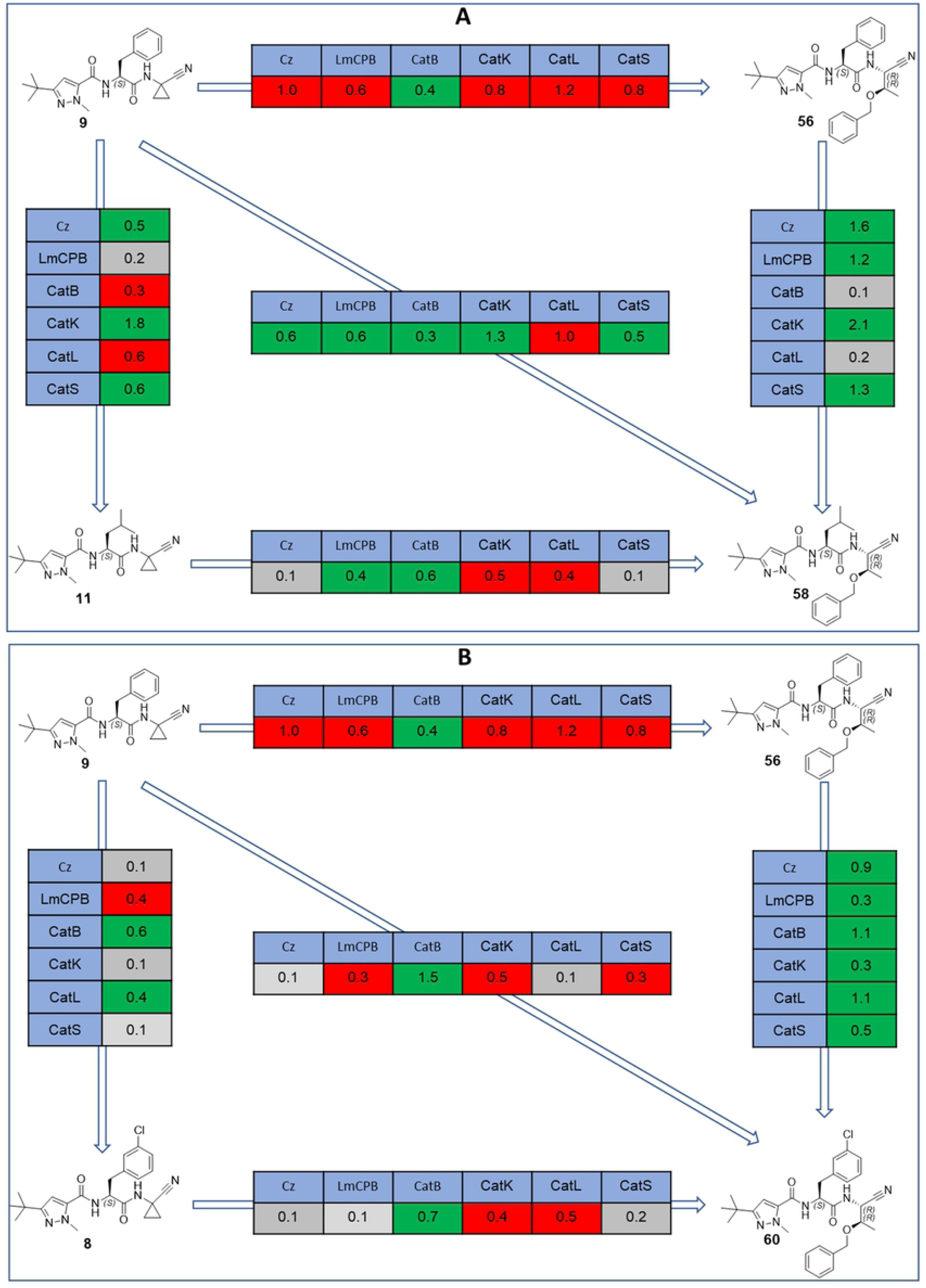
Non-additivity of SAR. Values are reported as differences in p*K*_i_ and are color-coded as red (negative), green (positive), grey (no significant difference, Δp*K*_i_< 0.2).

Pairwise plots for the selectivity towards Cz in relation to other human cathepsins are provided in Fig 11. It is not trivial to achieve a significant selectivity for Cz inhibitors (Δp*K*_i_ > 1.0) over mammalian CPs due to their high structural similarity of the active site. Undeniably, CatB has a different mode of binding due to the larger S2 and S3 pockets [31]. Compounds **14** and **67** displayed a significant selectivity toward CatL and CatS, respectively. The Cz selectivity in the case of compound **14** is driven by the S3-P3, while that of **67** second is driven by S1-P1 interaction. Additionally, the hydrophobic interaction in S1 and S1′ with P1 of compounds **50**, **54** and **60** resulted in a good selectivity over CatK. On the other hand, the results indicate how CatL, CatS and CatK inhibitors could be repurposed for the inhibition of protozoa CPs.

**Fig 11.**
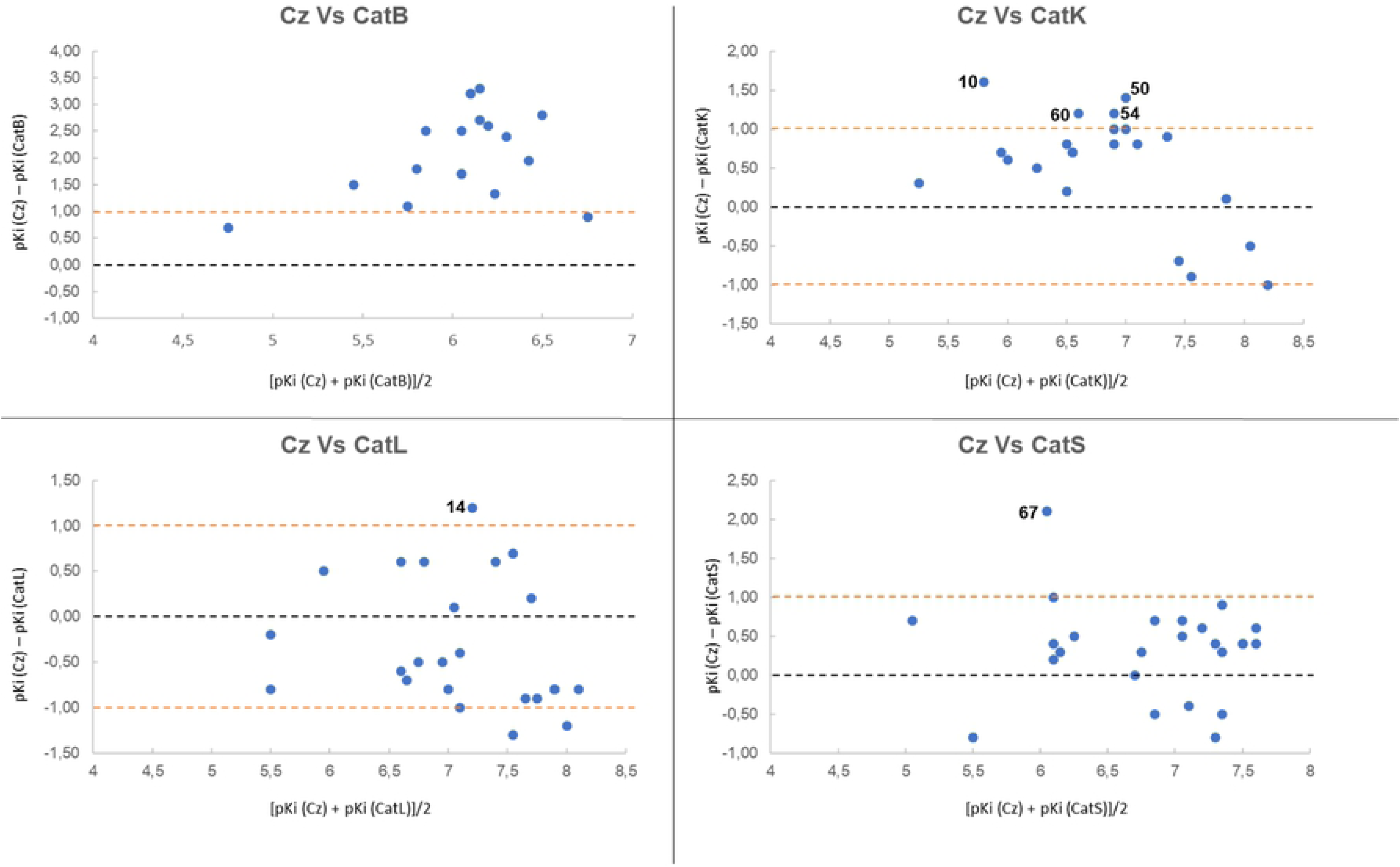
Selectivity pairwise plots. Values are given in p*K*_i_. X-axis represents the difference in p*K*_i_ for the same inhibitor for a pair of CPs. Y-axis represents the mean value of p*K*_i_ for the same inhibitor for a pair of CPs. Black dashed line highlights no selectivity. The magenta dashed line highlight a significant selectivity. Positive differences correspond to Cz p*K*_i_ values that are greater than those for CatB, CatK, CatL or CatS.

### Biological evaluation

All compounds synthesized were evaluated for their trypanocidal activity against the amastigote form of the Tulahuen *T. cruzi* strain, and the results are presented in Table 2. Three compounds (**52**, **57** and **60**) were equipotent with benznidazole as trypanocidal agents. In particular, compounds **52** and **60** are both low nanomolar Cz inhibitors and one-digit nanomolar inhibitors for CatL. Compound **57** had no affinity for any of the six CPs reported herein, which excluded the possibility that its mechanism of action is similar to compound **52** and **60**. Physiochemical properties (clogP, MM, TPSA, LogS) play an important role in drug design. As well, for potential trypanocidal agents, which had been designed as protozoan cysteine proteases inhibitors, physiochemical properties can influence their outcome. Therefore, we have included TPSA, calculated logP (ilogP), and LogS (Ali_LogS) in this discussion (Fig 12, Fig 13, Fig 14) [33].

**Fig 12.**
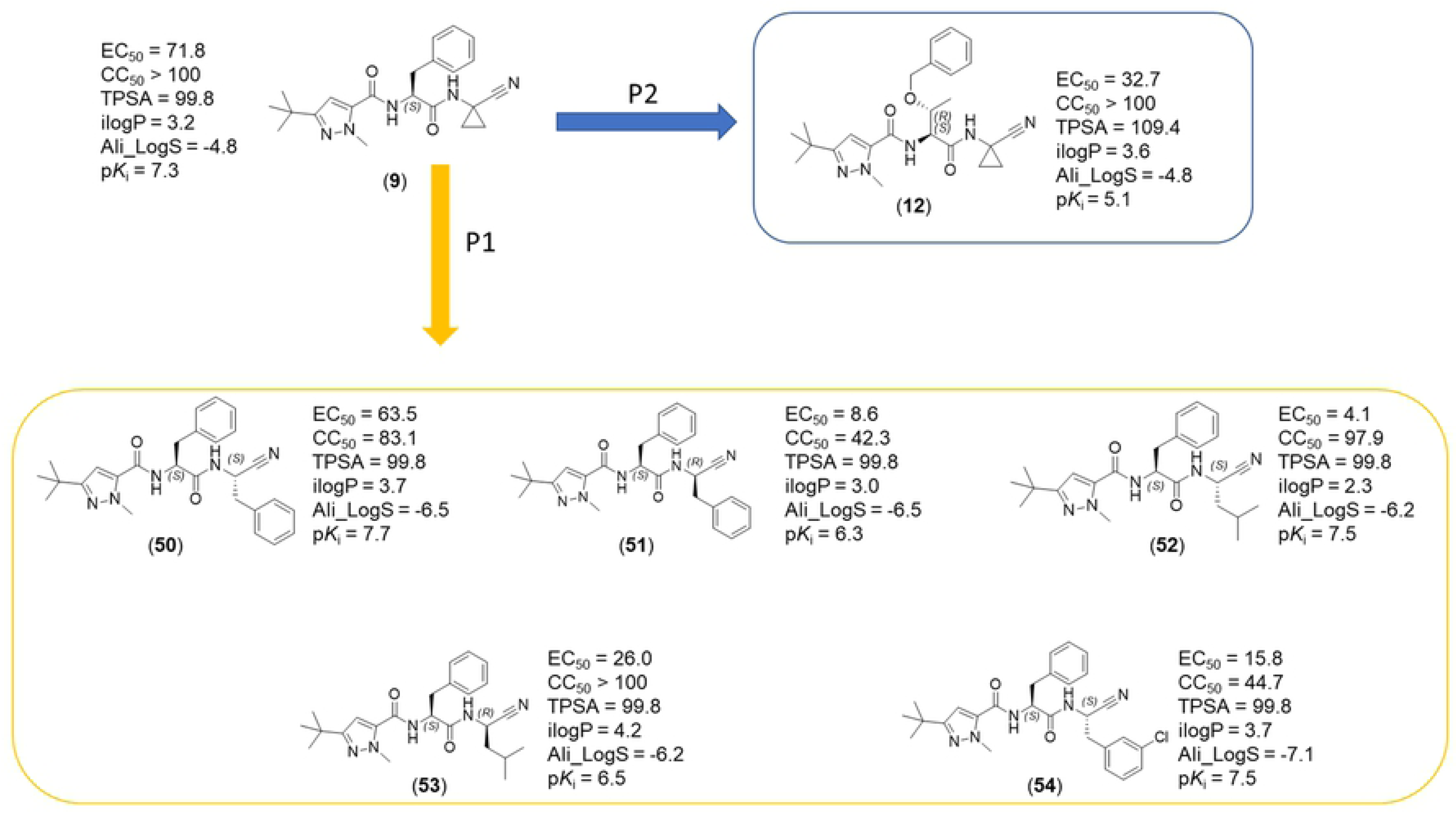
Schematic representation of physicochemical properties and SARs for trypanocidal activity. EC_50_ calculated for amastigote forms of *T. cruzi* (Tuhaluen strain). CC_50_ calculated for LLCMK2 strain (host cell). TPSA, ilogP, and Ali_Logs have been calculated with the swissADME on-line service [33].^1^ p*K*_i_ values are referring to Cz inhibition.

**Fig 3.**
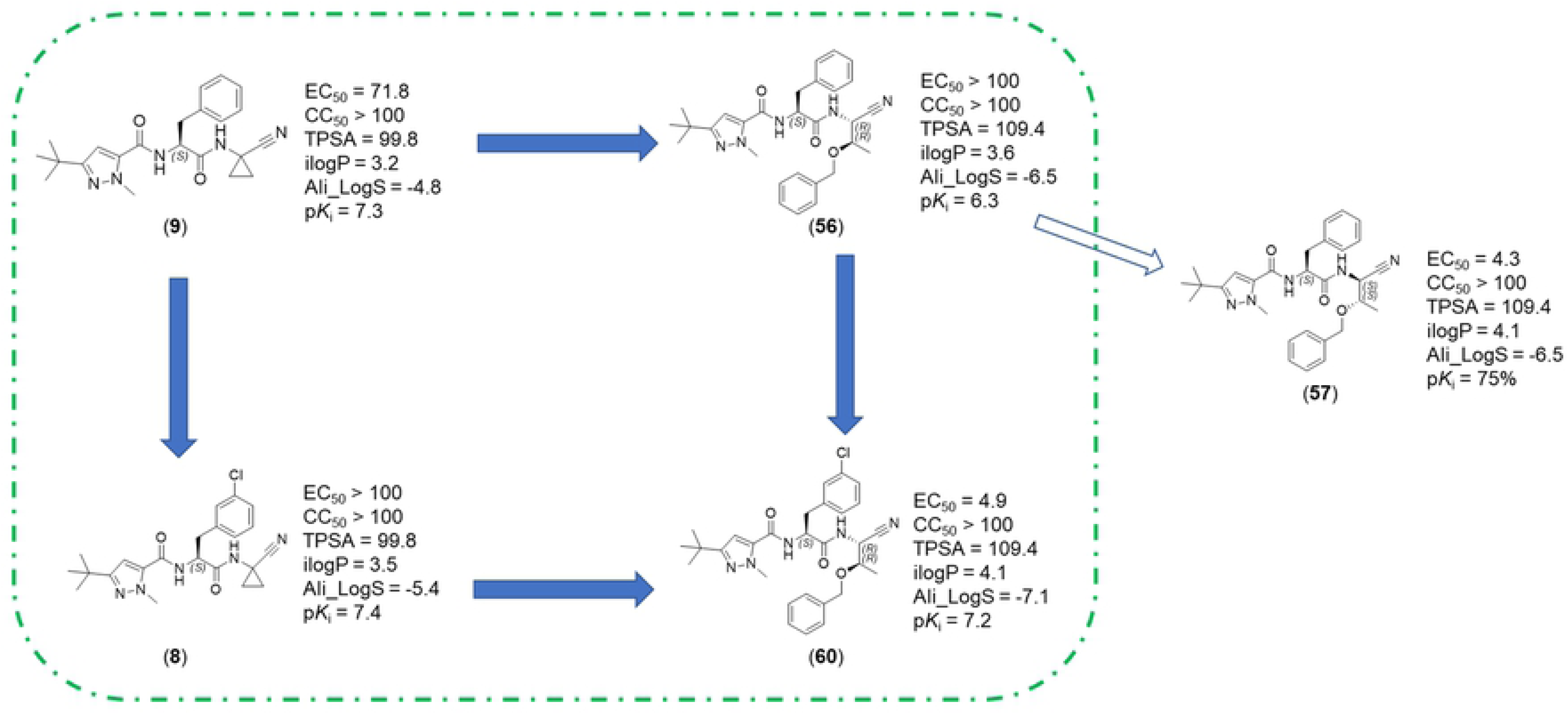
Schematic representation for non-additivity of SARs for trypanocidal activity. EC_50_ calculated for amastigote forms of *T.cruzi* (Tuhaluen strain). CC_50_ calculated for LLCMK2 strain (host cell). TPSA, ilogP, and Ali_Logs have been calculated with the swissADME on-line service.*^1^* p*K*_i_ values are referring to Cz inhibition.

**Fig 14.**
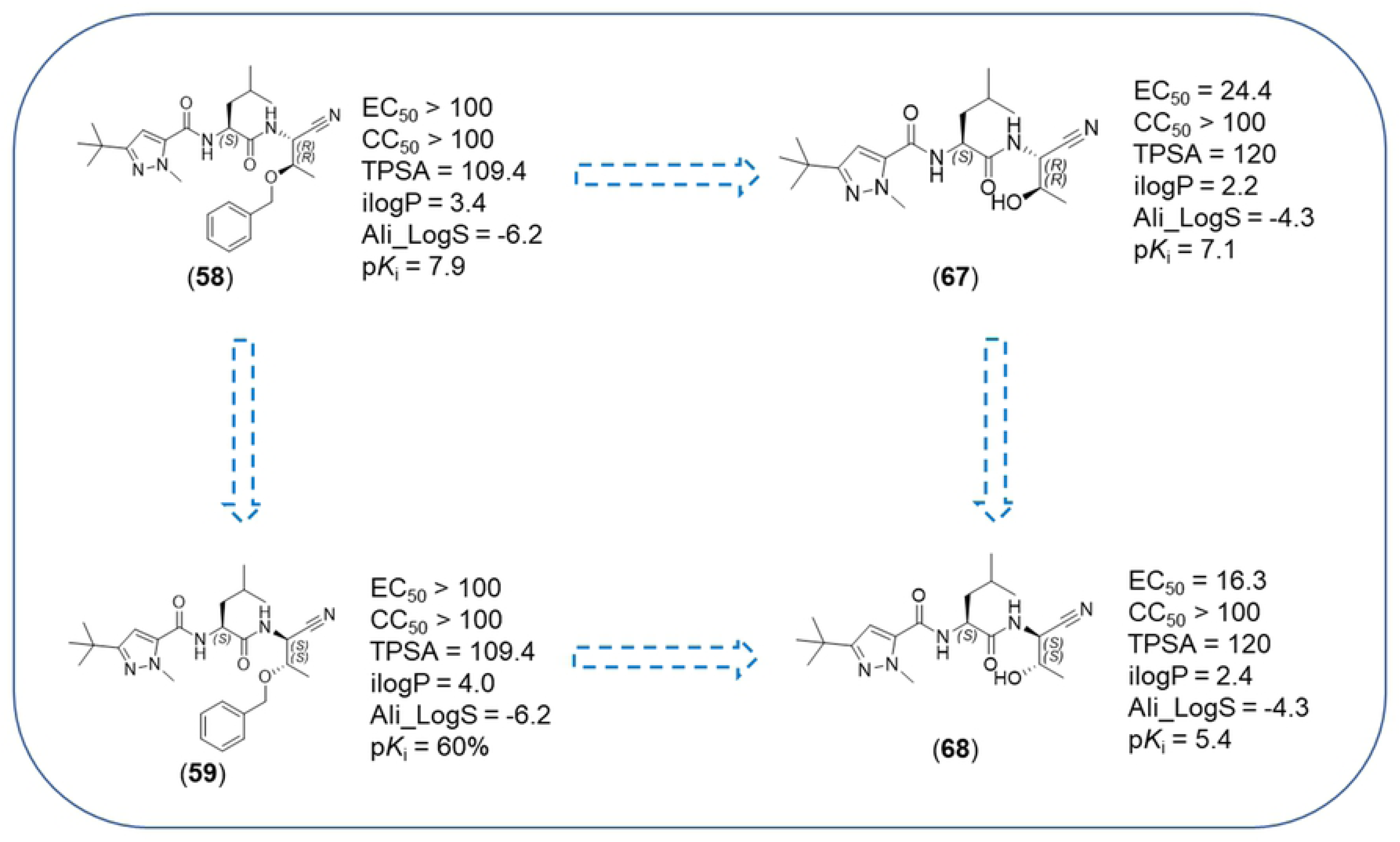
Schematic representation for non-additivity of SARs for compounds 58, 59, 67 and 68. EC_50_ calculated for amastigote forms of *T.cruzi* (Tuhaluen strain). CC_50_ calculated for LLCMK2 strain (host cell). TPSA, ilogP, and Ali_Logs have been calculated with the swissADME on-line service.*^1^* p*K*_i_ values are referring to Cz inhibition.

**Table 2.** Biological data for trypanocidal activity (EC_50_), cytotoxicity (CC_50_), and selective index (SI) for the series of dipeptidyl nitriles.

| Compounds | IC<sub>50</sub> (μM) | CC<sub>50</sub> (μM) | SI |
| --- | --- | --- | --- |
| **Bz** | 4.4 ± 0.47 | > 100 | > 23 |
| **51** | 8.6 ± 0.53 | 42.3 ± 3.94 | 4.9 |
| **52** | 4.1 ± 0.47 | 97.9 ± 6.52 | 24 |
| **57** | 4.3 ± 0.32 | > 100 | > 23 |
| **60** | 4.9 ± 0.45 | > 100 | > 20 |
| **9** | 72 ± 5.7 | > 100 | > 1.4 |
| **10** | 67.6 ± 8.22 | > 100 | > 1.5 |
| **12** | 32.7 ± 3.86 | > 100 | > 3.1 |
| **15** | 64.7 ± 4.32 | 73.6 ± 8.74 | 1.14 |
| **16** | 47.3 ± 3.26 | > 100 | > 2.1 |
| **50** | 63.3 ± 5.74 | 83.1 ± 6.85 | 1.31 |
| **53** | 26 ± 2.6 | > 100 | > 3.8 |
| **54** | 15.8 ± 1.43 | 44.7 ± 5.24 | 2.83 |
| **65** | 70.8 ± 8.79 | > 100 | > 1.4 |
| **66** | 30.0 ± 2.71 | > 100 | > 3.3 |
| **67** | 24.4 ± 2.36 | > 100 | > 4.1 |
| **68** | 16 ± 4.0 | > 100 | > 6.3 |
| **69** | 30.6 ± 3.11 | > 100 | > 3.3 |

Benznidazole was used as a positive control. DMSO was used to dissolve compounds and as a negative control. All data were obtained using at least two independent experiments. Compounds with EC<sub>50</sub> > 100 μM were considered to be not active, and they are not reported in this Table.

In general, the substitution of the P3 or P2 moieties from the prototype compound (**9**) did not result in any increment of potency, except in case of compound **12** (Fig 12). Substitution of Phe for (*S*,*R*)-Thr-O-Bn led to a modest trypanocidal activity. Modification in P1 strongly modulated the trypanocidal effect. If considering physicochemical properties, we observed a trend for this small compound set, insofar the reduction in lipophilicity corresponded to higher trypanocidal potency, as can be seen for compound **52** (EC_50_ = 4.1 µM; ilogP = 2.3) that presented 17.5 times more effective than initial molecule **9** (EC_50_ = 71.8 µM).

Single modification in P2 or P3 of compound **9** did not produce a beneficial effect for the trypanocidal activity. However, considering both modifications [**9**→**60**], compound **60** (EC_50_ = 4.93 µM) exhibited 14.5 times more effective than the compound **9** (EC_50_ = 71.8 µM) reaching values similar to Bz (gold standard drug to treated Chagaśdisease) exhibited a promising trypanocidal activity (Fig 13). In this case, the ilogP was the highest of the entire series and more than one log unit larger than for compound **54.** It is noteworthy that compound **57**, exhibiting the same values for TPSA and ilogP as **60**, had no affinity for CPs. In addition, the modifications in compound **57** (CC_50_ ≃ 124.7 µM) was not toxic to cells and showed to be the highest selectivity and lowest cytotoxicity in our assays. This guarantees an SI (selective index) ratio of over 20, as well as cysteine inhibitor compounds **60** and **52**, making them interesting targets for further *in vivo* testing against the acute form of chagas disease [34].

Compounds bearing a leucine moiety in P2 displayed a peculiar behavior (Fig 14). Indeed, debenzylation of the threonine moiety in P1 led to an increase in potency [**58**→**66**] and [**59**→**67**]. The trypanocidal potency for this set of compounds seemed not to correlate with Cz affinity; but, once again, active compounds had an ilogP value of less than 3.0.

Potential cytotoxicity of inhibitors was assessed with the LLCMK_2_ cell-based assay, and compounds were evaluated over three days using benznidazole as a control. Cytotoxicity at the highest concentration tested that did not lead to precipitation (250 μM) was low for the majority of test compounds. The most potent inhibitors of the amastigote *T*. *Cruzi* Tulahuen strains (**52, 57** and **60**) showed the same range of cytotoxicity when compared to benznidazole.

## Conclusion

In this study, we expanded our previous series of dipeptidyl nitrile inhibitors of Cz by leveraging the P1-S1/S1’ interaction. We studied how this interaction can influence affinity and selectivity for different CPs, also obtaining the inhibitory data for the whole series. Furthermore, 15 compounds had pEC_50_ above 4 against *T. cruzi* amastigote form, where three of them are equipotent with benznidazole as trypanocidal agents with SI (selective index) ratio of over 20, making them interesting targets for further *in vivo* testing against the acute form of chagas disease.

Based on the data obtained here and supported by our previous reports, the classic view of the small molecule entering the parasite and acting by inhibiting the critical function exerted by the cruzipain may be questioned. Many efforts have been made to introduce drugs with improved efficacy in order to cure Chagas disease, but not much is known about the mechanism of *T. cruzi* death. Even the mechanism of action of benznidazole is still poorly understood. The mere inhibition of intracellular cruzipain is likely insufficient to cause cell death, which implies several questions regarding the drug discovery approach and the current disease model, in particular whether the criteria for selecting cruzain as a target for therapeutic intervention are justified. Thus, also this study points to the common problem in the translation of the results from biochemical assays to the trypanocidal action. Our work also contributes to the understanding of subtle drug-target interactions and to the discovery of tailored trypanocidal agents equipotent to benznidazole and with the potential of further improvement.

## Supporting information Captions

**Table S1. Condition for enzymatic assays.**

**Table S2. Structures representation, number identification, nequimed number, pK_i_ values for CatB, CatK, CatL, CatS, Cz and LmCPB.**

**Table S3. Number identification, nequimed number, niological data for trypanocidal activity (EC_50_) and citotoxicity (CC_50_).**

**Figure S1. ^1^H NMR (500 MHz, DMSO_*d_6_*) of compound 7.**

**Figure S2. ^13^C NMR (125 MHz, DMSO_*d_6_*) of compound 7.**

**Figure S3. LC-MS report for compound 7.**

**Figure S4. ^1^H NMR (500 MHz, DMSO_*d_6_*) of compound 8.**

**Figure S5. ^13^C NMR (125 MHz, DMSO_*d_6_*) of compound 8.**

**Figure S6. LC-MS report for compound 8.**

**Figure S7. ^1^H NMR (200 MHz, CD_3_OD) of compound 10.**

**Figure S8. ^13^C NMR (50 MHz, CD_3_OD) of compound 10.**

**Figure S9. HPLC report of compound 10.**

**Figure S10. ^1^H NMR (200 MHz, CDCl_3_) of compound 12.**

**Figure S11** 1**^3^**C **NMR (50 MHz, CDCl_3_) of compound 12.**

**Figure S12. HPLC report of compound 12.**

**Figure S13. ^1^H NMR (500 MHz, DMSO_*d_6_*) of compound 13.**

**Figure S14. ^13^C NMR (125 MHz, DMSO_*d_6_*) of compound 13.**

**Figure S15. HPLC report for compound 13.**

**Figure S16. ^1^H NMR (400 MHz, DMSO_*d_6_*) of compound 14.**

**Figure S17. ^13^C NMR (100 MHz, DMSO_*d_6_*) of compound 14.**

**Figure S18. HPLC report ofr compound 14.**

**Figure S19. ^1^H NMR (500MHz, DMSO_*d_6_*) for compound 15.**

**Figure S20. ^13^C NMR (125 MHz, DMSO_*d_6_*) for compound 15.**

**Figure S21. HPLC report for compound 15.**

**Figure S22. ^1^H NMR (500MHz, DMSO_*d_6_*) for compound 16.**

**Figure S23. ^13^C NMR (125 MHz, DMSO_*d_6_*) for compound 16.**

**Figure S24. HPLC report for compound 16.**

**Figure S25. ^1^H NMR (200MHz, CD_3_OD) for compound 17.**

**Figure S26. ^13^C NMR (50 MHz, CD_3_OD) for compound 17.**

**Figure S27. HPLC report for compound 17.**

**Figure S28. ^1^H NMR (200MHz, CD_3_OD) of compound 18.**

**Figure S29. ^13^C NMR (50 MHz, CD_3_OD) of compound 18.**

**Figure S30. HPLC report for compound 18.**

**Figure S31. ^1^H NMR (200 MHz, CD_3_OD) of compound 19.**

**Figure S32. ^13^C NMR (50 MHz, CD_3_OD) of compound 19.**

**Figure S33. HPLC report for compound 19.**

**Figure S34. ^1^H NMR (400 MHz, CDCl_3_) of compound 50.**

**Figure S35. ^13^C NMR (100 MHz, CDCl_3_) of compound 50.**

**Figure S36. HPLC report of compound 50.**

**Figure S37. ^1^H NMR (200 MHz, CDCl_3_) of compound 51.**

**Figure S38. ^13^C NMR (50 MHz, CDCl_3_) for compound 51.**

**Figure S39. HPLC report of compound 51.**

**Figure S40. ^1^H NMR (200 MHz, CD_3_OD) of compound 52.**

**Figure S41. ^13^C NMR (50 MHz, CD_3_OD) of compound 52.**

**Figure S42. HPLC report of compound 52.**

**Figure S43. ^1^H NMR (200 MHz, CDCl_3_) of compound 53.**

**Figure S44. ^13^C NMR (50 MHz, CDCl_3_) of compound 53.**

**Figure S45. HPLC report of compound 53.**

**Figure S46. ^1^H NMR (200 MHz, CD_3_OD) of compound 54.**

**Figure S47. ^13^C NMR (50 MHz, CD_3_OD) of compound 54.**

**Figure S48. HPLC report of compound 54.**

**Figure S49. ^1^H NMR (400 MHz, CD_3_OD) of compound 55.**

**Figure S50. ^13^C NMR (100 MHz, CD_3_OD) of compound 55.**

**Figure S51. HPLC report of compound 55.**

**Figure S52. ^1^H NMR (200 MHz, CDCl_3_) for compound 56.**

**Figure S53. ^13^C NMR (50 MHz, CDCl_3_) for compound 56.**

**Figure S54. HPLC report of compound 56.**

**Figure S55. ^1^H NMR (200 MHz, CDCl_3_) of compound 57.**

**Figure S56. ^13^C NMR (50 MHz, CDCl_3_) of compound 57.**

**Figure S57. HPLC report of compound 57.**

**Figure S58. ^1^H NMR (400 MHz, DMSO_*d_6_*) of compound 58.**

**Figure S59. ^13^C NMR (100 MHz, DMSO_*d_6_*) of compound 58.**

**Figure S60. HPLC report of compound 58.**

**Figure S61. ^1^H NMR (400 MHz, CDCl_3_) of compound 59.**

**Figure S62. ^13^C NMR (100 MHz, CDCl_3_) of compound 59.**

**Figure S63. HPLC report of compound 59.**

**Figure S64. ^1^H NMR (400 MHz, CDCl_3_) of compound 60.**

**Figure S65. ^13^C NMR (100 MHz, CDCl_3_) of compound 60.**

**Figure S66. HPLC report of compound 60.**

**Figure S67. ^1^H NMR (400 MHz, CDCl_3_) of compound 65.**

**Figure S68. ^13^C NMR (100 MHz, CDCl_3_) of compound 65.**

**Figure S69. HPLC report of compound 65.**

**Figure S70. ^1^H NMR (400 MHz, CDCl_3_) of compound 66.**

**Figure S71. ^13^C NMR (100 MHz, CDCl_3_) of compound 66.**

**Figure S72. HPLC report of compound 66.**

**Figure S73. ^1^H NMR (400 MHz, CDCl_3_) of compound 67.**

**Figure S74. ^13^C NMR (100 MHz, CDCl_3_) of compound 67.**

**Figure S75. HPLC report of compound 67.**

**Figure S76. ^1^H NMR (400 MHz, CD_3_OD) of compound 68.**

**Figure S77. ^13^C NMR (100 MHz, CD_3_OD) of compound 68.**

**Figure S78. HPLC report of compound 68.**

**Figure S79. ^1^H NMR (400 MHz, CD_3_OD) of compound 69.**

**Figure S80. ^13^C NMR (100 MHz, CD_3_OD) of compound 69.**

**Figure S81. HPLC report of compound 69.**

**Figure S82. HPLC report with Diacel column of compound 50.**

**Figure S83. HPLC report Diacel column of compound 51.**

**Figure S84. HPLC report with Diacel column of a mixture of compounds 50 and 51.**

**Figure S85. HPLC report with Diacel column for compound 56.**

**Figure S86. HPLC report with Diacel column for compound 57.**

**Figure S87. HPLC report with Diacel column of a mixture of compound 56 and 57.**

**Figure S88. HPLC report with Diacel column for compound 65.**

**Figure S89. HPLC report with Diacel column for compound 66.**

**Figure S90. HPLC report with Diacel column of a mixture of compound 65 and 66.**

**Figure S91. Non linear kinetic plot for Neq0922 (58).**

**Figure S92. Regural residual plot for Neq0922 (58).**

**Figure S93. Plot of p*K*_i_ (Cz) vs. p*K*_i_ (LmCPB). A linear trendline fitted points.**

**Figure S94. Dose curve response for determination of CC_50_ (LLCMK2) and EC_50_ (*T. cruzi Tulahuen*).**

